# Obtaining deeper insights into microbiome diversity using a simple method to block host and non-targets in amplicon sequencing

**DOI:** 10.1101/2020.10.05.322305

**Authors:** Teresa Mayer, Alfredo Mari, Juliana Almario, Mariana Murillo-Roos, Muhammad Abdullah, Nina Dombrowski, Stephane Hacquard, Eric M. Kemen, Matthew T. Agler

## Abstract

Microbiome profiling is revolutionizing our understanding of biological mechanisms such as metaorganismal (host+microbiome) assembly, functions and adaptation. Amplicon sequencing of multiple conserved, phylogenetically informative loci is an instrumental tool for characterization of the highly diverse microbiomes of natural systems. Investigations in many study systems are hindered by loss of essential sequencing depth due to amplification of non-target DNA from hosts or overabundant microorganisms. This issue requires urgent attention to address ecologically relevant problems using high throughput, high resolution microbial profiling. Here, we introduce a simple, low cost and highly flexible method using standard oligonucleotides (“blocking oligos”) to block amplification of non-targets and an R package to aid in their design. They can be dropped into practically any two-step amplicon sequencing library preparation pipeline. We apply them in leaves, a system presenting exceptional challenges with host and non-target microbial amplification. Blocking oligos designed for use in eight target loci reduce undesirable amplification of host and non-target microbial DNA by up to 90%. In addition, 16S and 18S “universal” plant blocking oligos efficiently block most plant hosts, leading to increased microbial alpha diversity discovery without biasing beta diversity measurements. By blocking only chloroplast 16S amplification, we show that blocking oligos do not compromise quantitative microbial load information inherent to plant-associated amplicon sequencing data. Using these tools, we generated a near-complete survey of the *Arabidopsis thaliana* leaf microbiome based on diversity data from eight loci and discuss complementarity of commonly used amplicon sequencing regions for describing leaf microbiota. The blocking oligo approach has potential to make new questions in a variety of study systems more tractable by making amplicon sequencing more targeted, leading to deeper, systems-based insights into microbial discovery.

## Introduction

A revolution in biology is currently underway as our understanding of various systems is brought into the context of the structures and roles of symbiotic microbial consortia. This transformation is the result of increasing research to characterize the microbiota associated with various abiotic or biotic systems. For example, important roles of microbial communities have been revealed in systems as diverse as biotechnological transformations^1^ and plant and animal health and fitness^2–5^. To do so, many studies rely on microbiota profiles generated by amplicon sequencing of phylogenetically informative genomic loci. These profiles are then linked to specific experimental parameters, host phenotypes or performance measurements^6^.

Microbiomes often include species from all kingdoms of life. These cohabiting members interact with the environment and influence one another via direct associations ^7^ or indirectly via a host ^8^. To resolve these interactions and model microbial community dynamics, robust systems approaches are needed^9^ with integration of diversity beyond bacteria^10^. Such approaches have revealed, for example, keystone species that participate heavily in inter-kingdom interactions in phyllosphere microbial communities^11^ and in ocean samples^12^ and which thereby underlie microbial community structures. Whatever the system, robust approaches to pinpoint important microbes in community surveys require broad and deep coverage of diversity in a high-throughput manner. Additionally, quantitative abundance data is needed to accurately infer inter-microbial interactions.

Researchers use many technologies and pipelines to generate and sequence amplicon libraries. A major problem affecting broad-diversity amplicon sequencing pipelines is that “universal” amplification primers amplify DNA from non-target or overabundant organisms (e.g., hosts^13,14^, resident sporulating microorganisms^11^ or endosymbionts^15^), reducing effective sequencing depth and obscuring microbial diversity. Methods commonly used to address this problem include peptide nucleic acid (PNA) “clamps”^16^ or oligonucleotides modified with a C3 spacer^17^, which both arrest amplification of non-target amplicons. These, however, can be costly to design and implement, especially when the needs of researchers are constantly changing. Additionally, non-target (e.g., host) abundance information can provide quantitative insights into microbial load^18^, and none of these methods are designed to retain this quality. We employed amplicon sequencing to generate microbial diversity data from multiple loci from 16S and 18S rRNA genes (bacteria and eukaryotes, respectively) as well as the internal transcribed spacer (ITS) of fungi and oomycetes. We target microbial diversity in plant leaves, a challenging system where amplification of non-target and occasionally sporulating microbiota is extensive, resulting in large amounts of wasted data. To address this major barrier, we introduce a new method that uses a pair of standard oligonucleotides, making it cost-efficient and flexible. Additionally, they can be dropped into almost any library preparation pipeline. Indeed, this method, which we first applied in 2016^11^, has since been used successfully in multiple studies^19,20^ but its applicability and accuracy has not yet been broadly tested. Here, we extend the approach to 8 loci in the 16S, 18S, ITS1 and ITS2 regions and demonstrate it is effective in blocking most host plant species and a non-target microorganism without biasing diversity results. We also show that in plants, increasing read depth and diversity discovery with blocking oligos is compatible with deriving quantitative bacterial load information from 16S data. This simple solution enables rapid and nearly complete characterization of hyperdiverse microbiomes in difficult systems and increases diversity discovery, broadening the applicability and impact of amplicon sequencing experiments. Finally, we provide an “R” package with three simple functions to rapidly and easily design oligos to block amplification of any specific DNA template.

## Results

### Blocking oligos reduce non-target amplification by “universal” primers

Host or other non-target amplicons are not useful to assess microbial diversity and are therefore often discarded, wasting sequencing depth. Therefore, we developed “blocking oligos” to reduce amplification of non-target DNA. Blocking oligos are standard oligonucleotides whose binding site is nested inside the binding site of “universal” primers for a locus of interest and are highly specific for a non-target organism **(Fig. 1** and **Fig. S1)**. During the first PCR step (blocking cycles), their nested binding location physically blocks the non-target elongation by the polymerase at the “universal” primer site, resulting in only short non-target amplicons. In the second PCR step (extension cycles), concatenated primers are used to add indices and Illumina sequencing adapters. Since the concatenated primer binding site is not present on non-target products, they are not amplified, and the sequencing library becomes enriched with target amplicons (**Fig. 1**). They can be dropped into the first step of any standard two-step amplicon library preparation pipeline.

**Figure 1.**
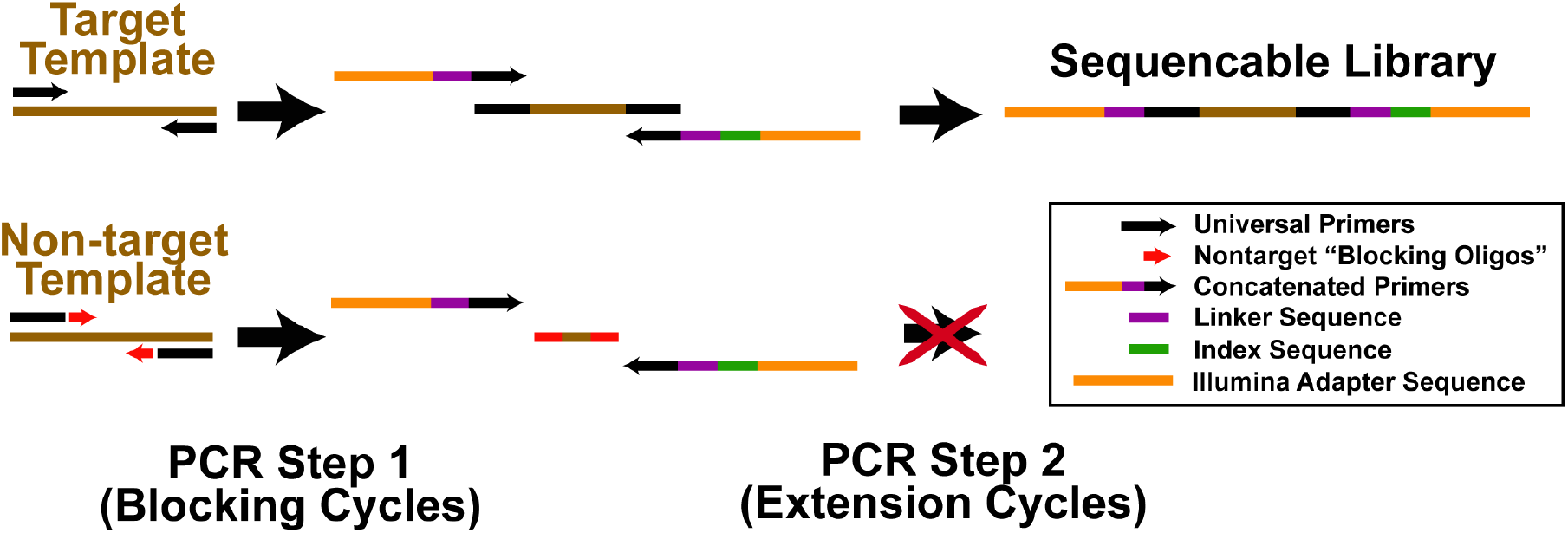
A strategy to reduce non-target amplification in amplicon sequencing pipelines. A 2-step amplification approach is used in which universal primers first amplify genomic templates, then indices and adapters are added in a second step. To prevent amplification of non-target templates, blocking oligos complementary to non-target genomic templates are employed in the first “blocking cycles” PCR step, resulting in short amplicons that cannot be amplified with concatenated primers in the second “extension cycles” step. Without addition of Illumina adapter sequences, these PCR products are not sequenced.

We previously designed blocking oligos to reduce amplification of plant chloroplast (16S V3-V4 rRNA), mitochondria (16S V5-V7 rRNA) and plant ITS (fungal and oomycete ITS regions 1 and 2) (**Table 1**)^11^. Here, we thoroughly tested them by checking how much they reduced host amplification compared to a “standard” library preparation without blocking, and whether they biased beta diversity estimates. We used a mock community that simulated a host associated microbiome (95% *A. thaliana* / 5% microbial genomic DNA). With blocking oligos, useful read depth was largely recovered by eliminating 60 - 90% of non-target contamination in bacterial 16S datasets and nearly all of the small amount of contamination in fungal ITS data (**Fig 2A and 2B**). Blocking oligos were slightly more effective in even (target microbes all in equal abundance) than in uneven communities (target microbes in unequal abundance), but much of this difference was either caused by poor taxonomic annotation or universal primer bias (see **Supporting Notes**). Importantly, in all three kingdoms, replicates that only differed in the use of blocking oligos were very similar to the expected distribution, demonstrating that blocking oligos do not change the recovered taxa distribution (shown at the genus level in **Fig 2A, Fig S2, Fig S3** and at the order level in **Fig S4**). Variations of the library preparation protocol had little or no effect on the results (see **Supporting Note** for additional details on testing the library preparation protocol).

**Figure 2.**
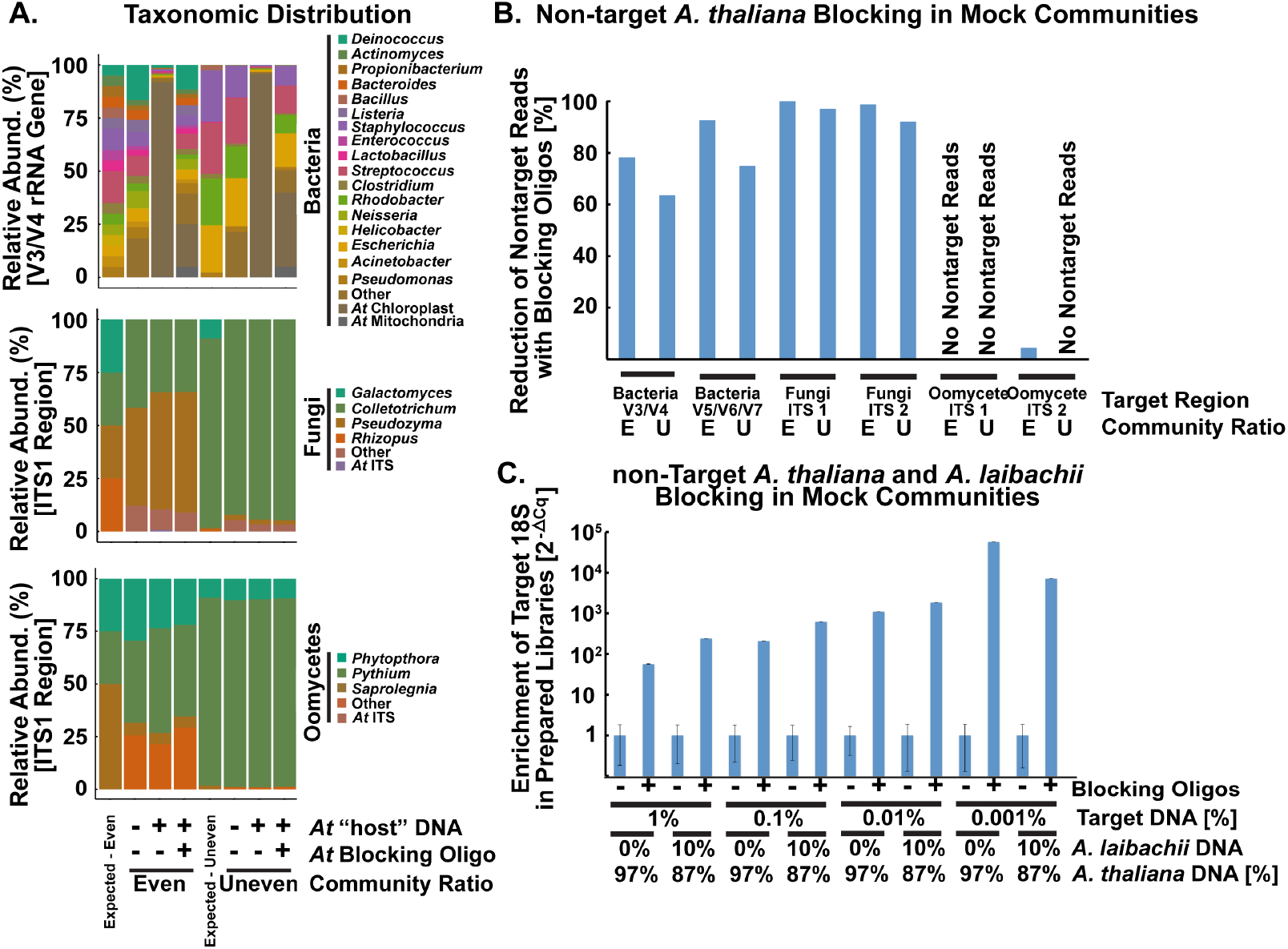
Reproducible and accurate characterization of mock communities of bacteria, fungi, and oomycetes by amplicon sequencing. (A) Observed taxa at the genus level in sequenced mock communities closely matched expected communities. The taxa “Other” is primarily non-target amplification from *A. thaliana* “host” DNA that was added to test blocking oligomers which prevent “host” DNA amplification. “NA” indicates a sample where sequencing depth was too low after subsampling to be included. (B) Near-complete reduction of amplification of *A. thaliana* “host” non-target plastid 16S or ITS by employing blocking oligos in preparation of mock community libraries. “E” and “U” refer to even and uneven communities, respectively. (C) Relative increase of target (*Saccharomyces* sp.) 18S V4-V5 region amplicons (qPCR 2^-ΔCq^ values relative to measurement without blocking oligomers) in mock community libraries prepared with blocking oligomers to reduce *A. thaliana* and *A. laibachii* non-target amplification.

**Table 1:**
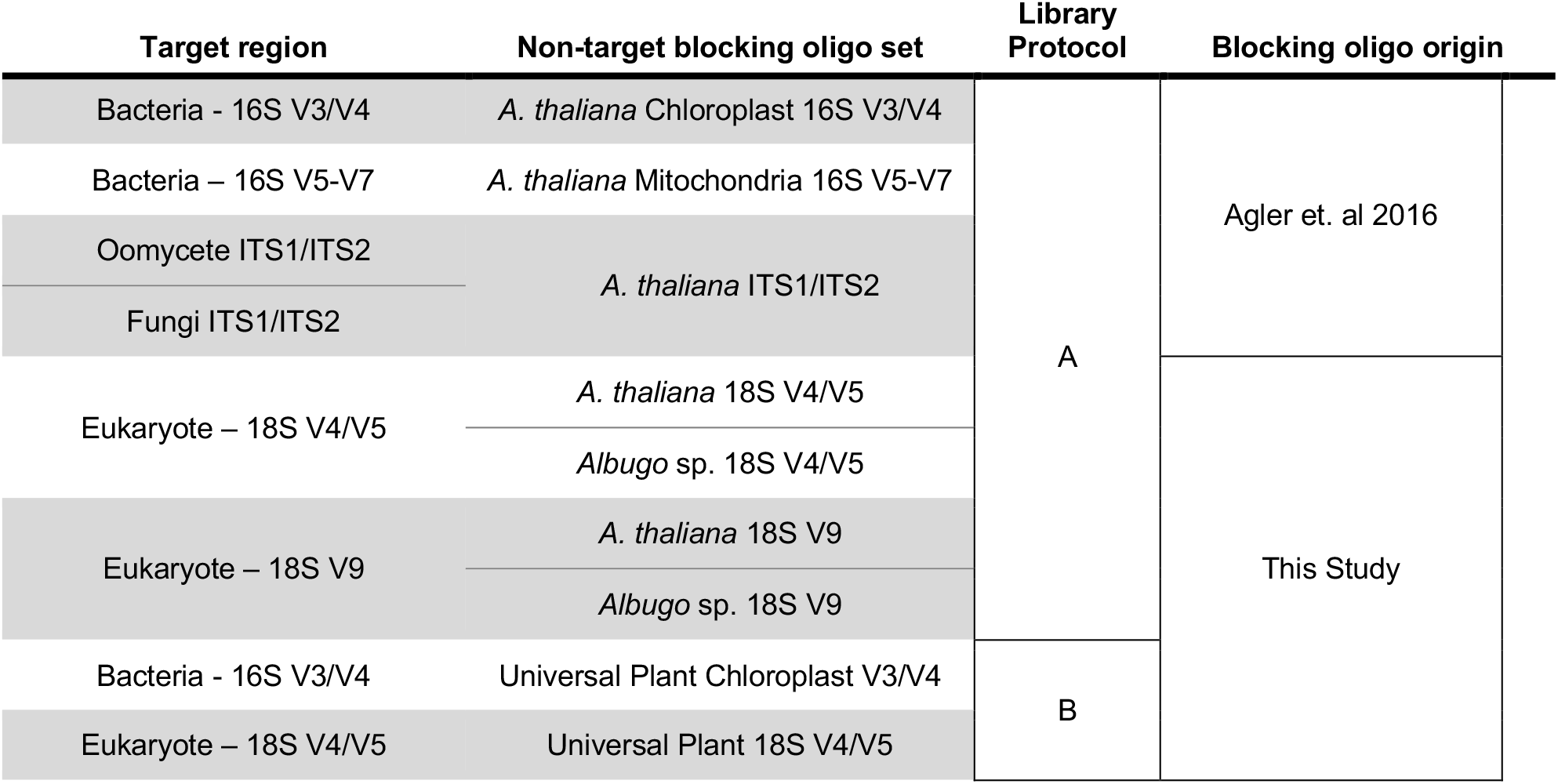
Overview of the loci and the non-target regions for which blocking oligos were designed previously and for this study. The blocking oligo approach of Agler et. al (2016) was extended here to 8 loci to characterize bacteria, fungi, oomycetes and other eukaryotic microbiota while avoiding non-target *A. thaliana* and *Albugo* sp. amplification. Libraries were prepared and sequenced with “Protocol A”, similar to Agler et. al (2016). Single sets of blocking oligos that block amplification of DNA of most plant hosts were designed for the 16S and 18S regions. An alternative protocol B was used for sequencing and library preparation. All primer sequences are available in Files S1a and S1b.

Sequencing 18S rRNA gene libraries in addition to ITS (**Table S1**) should recover more leaf eukaryotic microbial diversity^21^. This diversity would be obscured, however, by the host and occasionally by sporulating pathogens like *Albugo laibachii* that are efficiently amplified by universal 18S primers. Therefore, we designed and tested blocking oligos to overcome non-target amplification of both *A. laibachii* and *A. thaliana* in the 18S region (**Table 1)**. For testing, we generated mock genomic DNA templates (**Table S2**) containing bacterial (*Bacillus* sp.), *A. thaliana, A. laibachii* and target (*S. cerevisiae*) genomic DNA. We then prepared 18S V4-V5 region amplicon libraries from the template samples with or without *A. thaliana* and *A. laibachii* blocking oligos in the first PCR step. Finally, we quantified the levels of target (*S. cerevisiae*) amplicons in the prepared libraries using qPCR. Indeed, blocking both nontargets increased target levels between ~57x (1% target template) and ~57,000x (0.001% target template) (**Fig 2C**), demonstrating that both host and non-target microbial taxa can be efficiently simultaneously blocked.

### “Universal” plant blocking oligos enable profiling of microbiota in many host species

*A. thaliana* blocking oligos are not effective against every plant host, so users would need to design and test new oligos for their purposes^22^. Thus, we expanded to multiple hosts by designing a set of oligos to block amplification of chloroplast 16S rRNA and plant 18S rRNA genes using a highly diverse set of plant species (see **Table S3**). Candidates were first tested for specificity to plants by amplifying DNA from 21 plant species (**Table S4**) and mixed bacterial DNA and then visualizing bands on a gel. We selected primer sets that amplified most plants but avoided amplification of bacteria or fungi (**Fig S5**). Next, we tested the oligos by preparing sequencing libraires (Protocol B, see supplementary notes for comparison with Protocol A) with DNA templates from leaves of plant species representing five orders spanning monocots and dicots (*Amaranthus spec., Arabidopsis thaliana, Bromus erectus, Lotus corniculatus* and *Plantago lanceolata*) (see **Table S5**). Although *A. thaliana* blocking oligos were very efficient in mock leaf microbiomes (**Fig 2**), in real leaf samples they only sometimes helped recover higher microbial diversity (**Fig S6D/E and S7D/E**). Microbial loads on leaves are typically very low^20^, so we reasoned that more blocking cycles may be required in real leaves. Therefore, for testing universal 16S blocking oligos we compared 10 vs. 15 blocking cycles.

With 10 blocking cycles, 1-25% of target (non-chloroplast) reads were recovered from *Arabidopsis thaliana*, *Bromus erectus* and *Lotus corniculatus* (**Fig 3A**, the other two species had no usable reads with 10 cycles). 15 blocking cycles increased the amount of retrieved target reads by at least 2.5-fold compared to 10 cycles (**Fig 3A**), increasing the fraction of useful reads 8-16x compared to without blocking oligos in all five plant species (**Fig 3A** and **Fig S8**). 18S blocking oligos were only tested with 10 blocking cycles but in four plant species we observed an increase from < 5% target (non-plant) reads without blocking oligos to up to 57% target reads with blocking oligos (**Fig S9**). Next, we again checked whether blocking oligos bias recovered beta diversity (differences between samples). In the 16S, we observed no significant effects on leaf samples (**Fig 3B**), a microbial community standard (**Fig S10**), nor in three different samples with soil DNA as template (**Fig S11**). We only tested the 18S oligos on leaf samples and observed that they resulted in recovery of more diverse communities. However, without further testing using mock communities we cannot say to which extent, if any, the 18S oligos introduce bias to the measurements. Overall, all universal blocking oligos can be used with practically any sample to increase useful data recovery and 16S blocking oligos do this without biasing beta diversity patterns.

**Figure 3:**
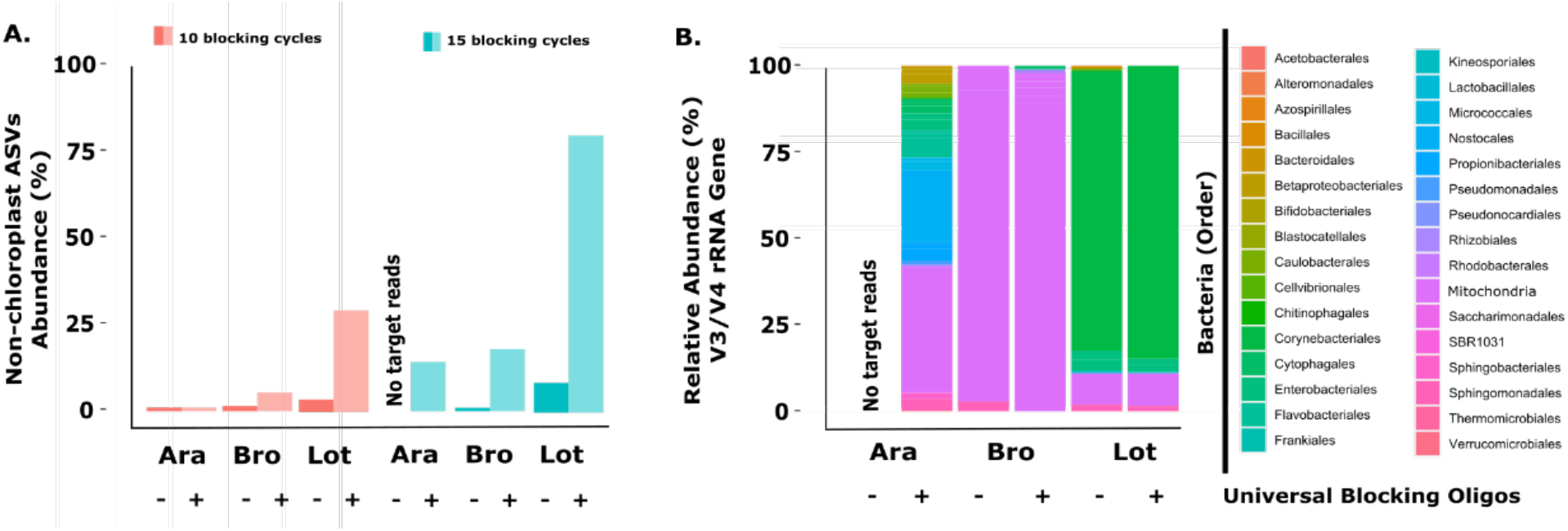
Universal blocking oligos successfully block undesired reads whilst remaining specific for recovered bacterial species and therefor increasing sequencing depth. (A) Percentage of reads assigned to ASVs other than chloroplast (non-chloroplast ASVs) with 10 vs 15 blocking cycles. The use of blocking oligos leads to higher recovery of bacterial ASVs. When the number of blocking cycles is increased, the fraction of blocked ASVs increases as well. (B) Taxonomic distribution in samples of different plants species with and without blocking oligos (15 blocking cycles). The use of universal blocking oligos does not significantly change the identity of retrieved ASVs. Results for other plant species and mock communities are shown in Fig S6 and Fig S7.

### “Universal” plant blocking oligos increase recovered alpha diversity

When the majority of reads retrieved from amplicon sequencing are non-target, the effective sequencing depth is drastically decreased. Thus, an important question is whether this actually obscures the microbial diversity recovered and whether blocking oligos allow recovery of higher alpha diversity. We checked Shannon and Simpson (observed species richness and diversity, respectively) and ACE and Chao1 (which estimate total species richness) alpha diversity of bacterial 16S data from the three naturally grown plant species amplified with 10 or 15 blocking cycles with and without universal blocking oligos. With 10 blocking cycles, observed richness and diversity were marginally higher (**Fig 4A and 4B**) and estimates of total species richness were unchanged (**Fig 4C and 4D**) with blocking. Blocking for 15 cycles, on the other hand, resulted in significantly higher observed and estimated richness and diversity (p<0.01 for Shannon and Simpson and p=0.11 and p=0.09 for Chao1 and ACE, respectively) (Fig 4). The difference between 10 and 15 cycles is again most likely due to low bacterial loads in real leaf samples (**Fig S6D/E and S7D/E**). Thus, >10 blocking cycles are recommended to consistently recover complete diversity.

**Figure 4:**
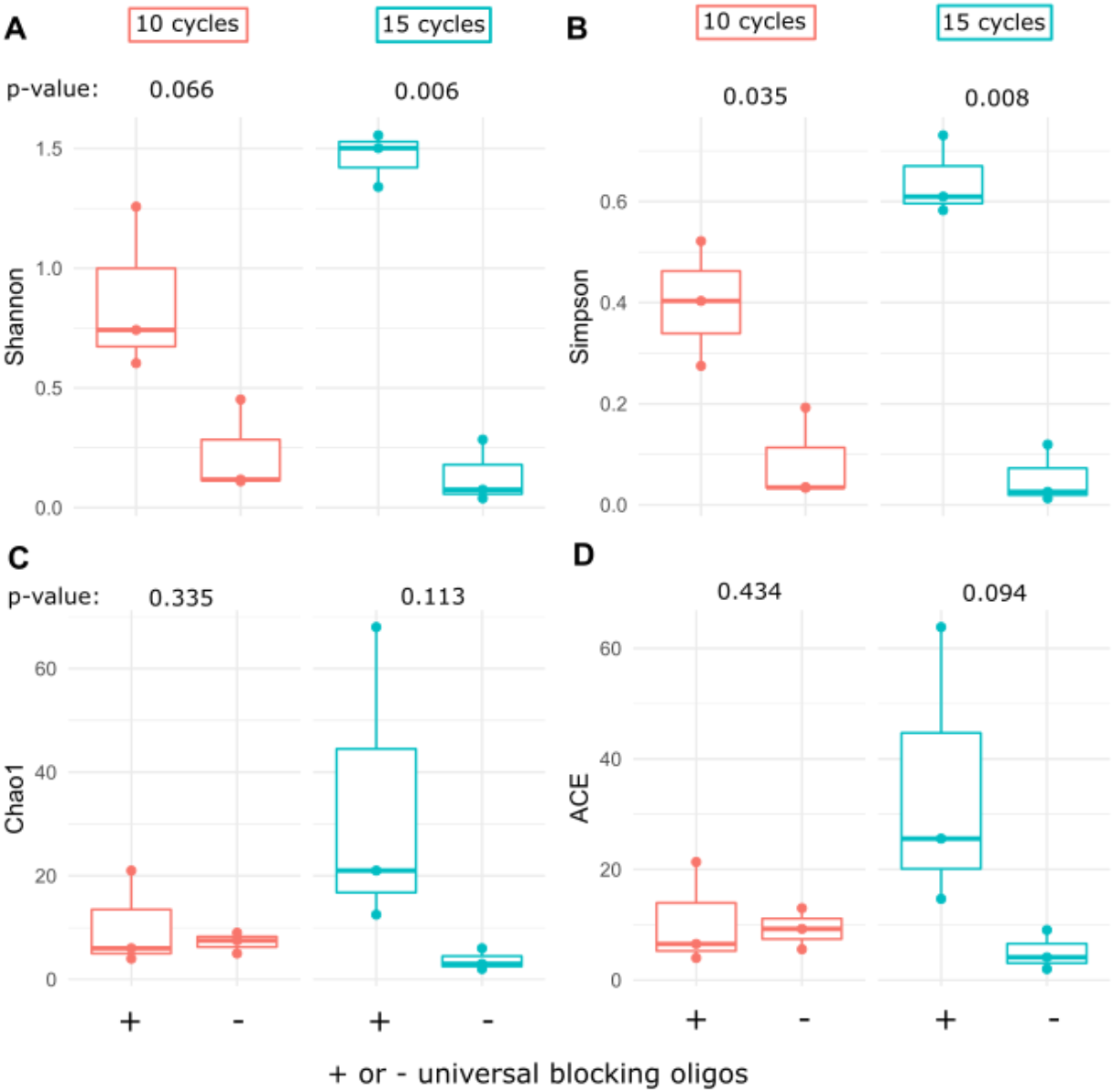
The use of blocking oligos increases the bacterial alpha diversity recovered. Comparison of alpha diversity measures between samples with 10 or 15 blocking cycles. We calculated the alpha diversity indices Shannon (A), Simpson (B), Chao1 (C) and ACE (D). Shannon and Simpson diversity indices combine richness and diversity (they measure both the number of species as well as the inequality between species abundances), whereas Chao1 and ACE estimate the total species richness. The use of blocking oligos for 10 cycles showed an increase in only Shannon and Simpson alpha diversity indices. However, when blocking for 15 cycles all indices show an increase in alpha diversity. p-values are calculated with a one-sided paired t-test.

### Leaf bacterial loads can be estimated using 16S amplicon data

One limitation of amplicon sequencing is that it is compositional, such that the quantitative bacterial load information is lost. Recently it has been demonstrated that “non-target” host to target bacterial ratios can be used to roughly estimate bacterial loads in plant samples^18^. Losing this information would be a downside of implementing blocking oligos. We observed that after blocking chloroplast amplification with the universal blocking oligos, host mitochondrial reads still made up a significant part of the data (**Fig 3**).

Therefore, we checked whether quantitative load information is still contained in the data generated with chloroplast blocking oligos. We tested this using the same plant species as before, which we grew axenically, harvested DNA, then combined with specific amounts of a bacterial DNA mix (Zymo Research Europe, Gmbh). We then estimated the fraction of bacterial 16S sequences recovered (after filtering remaining chloroplast reads – **Fig. 5**).

**Figure 5:**
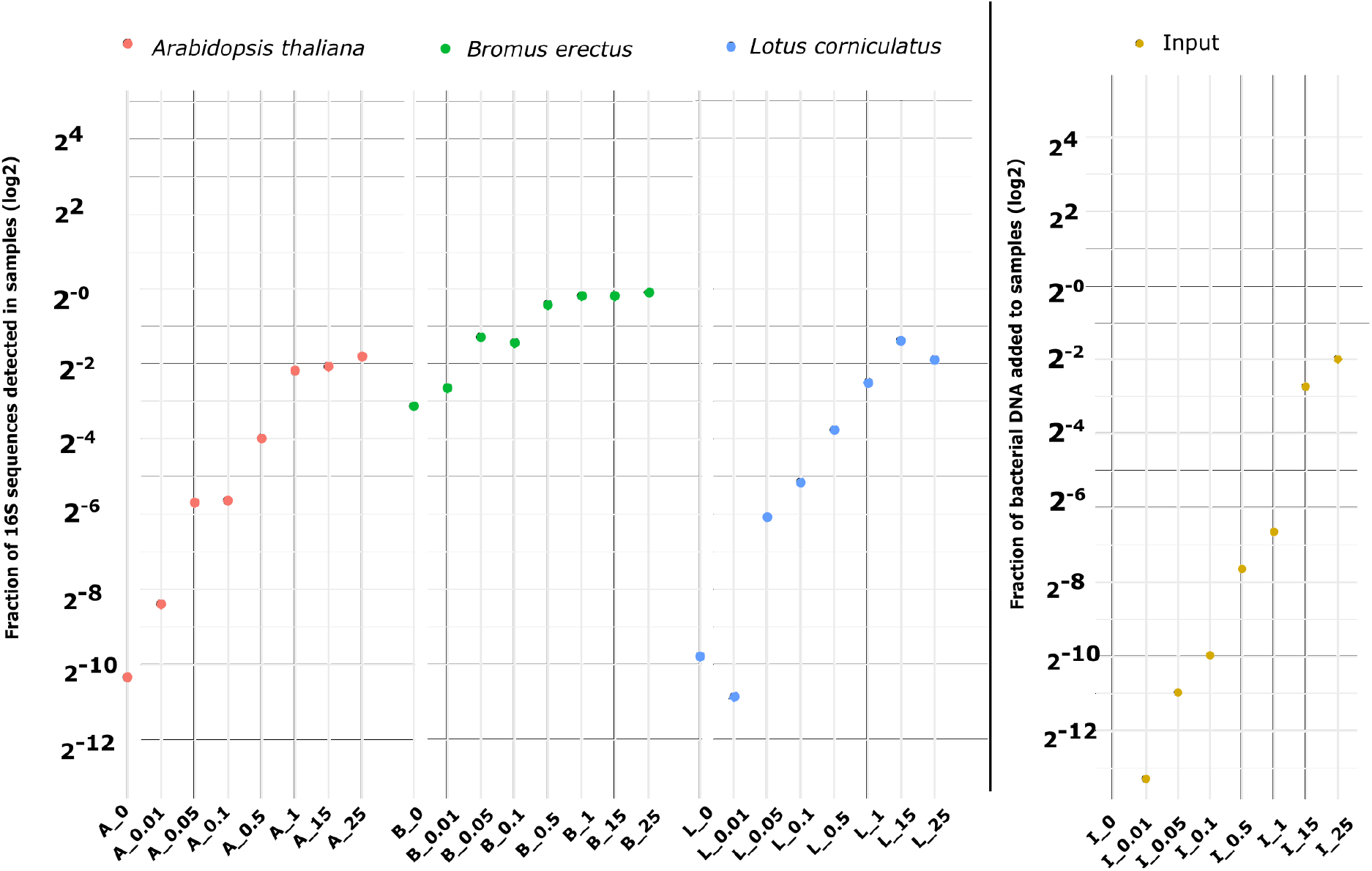
The fraction of bacterial reads can be used to gain quantitative microbial load information. The fraction of bacterial reads in sequenced amplicon libraries increases with as the load of mixed bacterial gDNA increases in the template. The axenic plant gDNA used in the mixes *were A. thaliana* (red), *B. erectus* (green) and *L. corniculatus* (blue). The fraction of bacterial gDNA compared to plant gDNA is shown in yellow (“Input”).

Up to a 1% fraction of bacterial DNA in the template, we observed a nearly linear increase in the fraction of sequenced reads assigned to bacteria (**Fig 5**). For a given load, the fraction of bacterial reads was similar for *A. thaliana* and *L. corniculatus* and was higher for *B. erectus*. Therefore, we conclude that within samples of the same species blocking oligos not only increase recovered bacterial diversity but can be applied so that quantitative bacterial load information is maintained.

### An expanded multi-kingdom view of leaf microbial diversity

We tested using the blocking oligo system to generate as broad of a microbial diversity profile as possible from leaves. We amplified and sequenced the 8 target loci in 12 wild *A. thaliana* leaf samples, including leaves with sporulating *A. laibachii* infections, and employed the *A. thaliana* and *A. laibachii* specific blocking oligos. Then we analyzed the diversity insights gained with this broad approach. The addition of the 18S rRNA gene primers broadly targeting eukaryotes increased diversity recovery by nearly 50% compared to ITS primers alone (observed genera, **Fig 6**). This included red and green algae, cercozoa and amoebozoa and even suggested signs of metazoa like insects and helminthes (**Fig 6** and **File S2**). The fungal and oomycete ITS datasets complemented the broader 18S data with more specificity in those groups – together, these two accounted for 44% of observed eukaryotic genera (**Fig 6a**). Prokaryote datasets further demonstrate complementarity for primer sets targeting the same groups of microbes (**Fig 6B**). Here, 42% of observed genera were discovered by both primer sets, with complementary diversity discovery especially in the phyla *Cyanobacteria* (V3-V4 dataset) and *Firmicutes* (V5-V7 dataset). Thus, blocking of over-abundant host and microbial amplicons allows deep diversity characterization in leaves using multiple loci.

**FIGURE 6.**
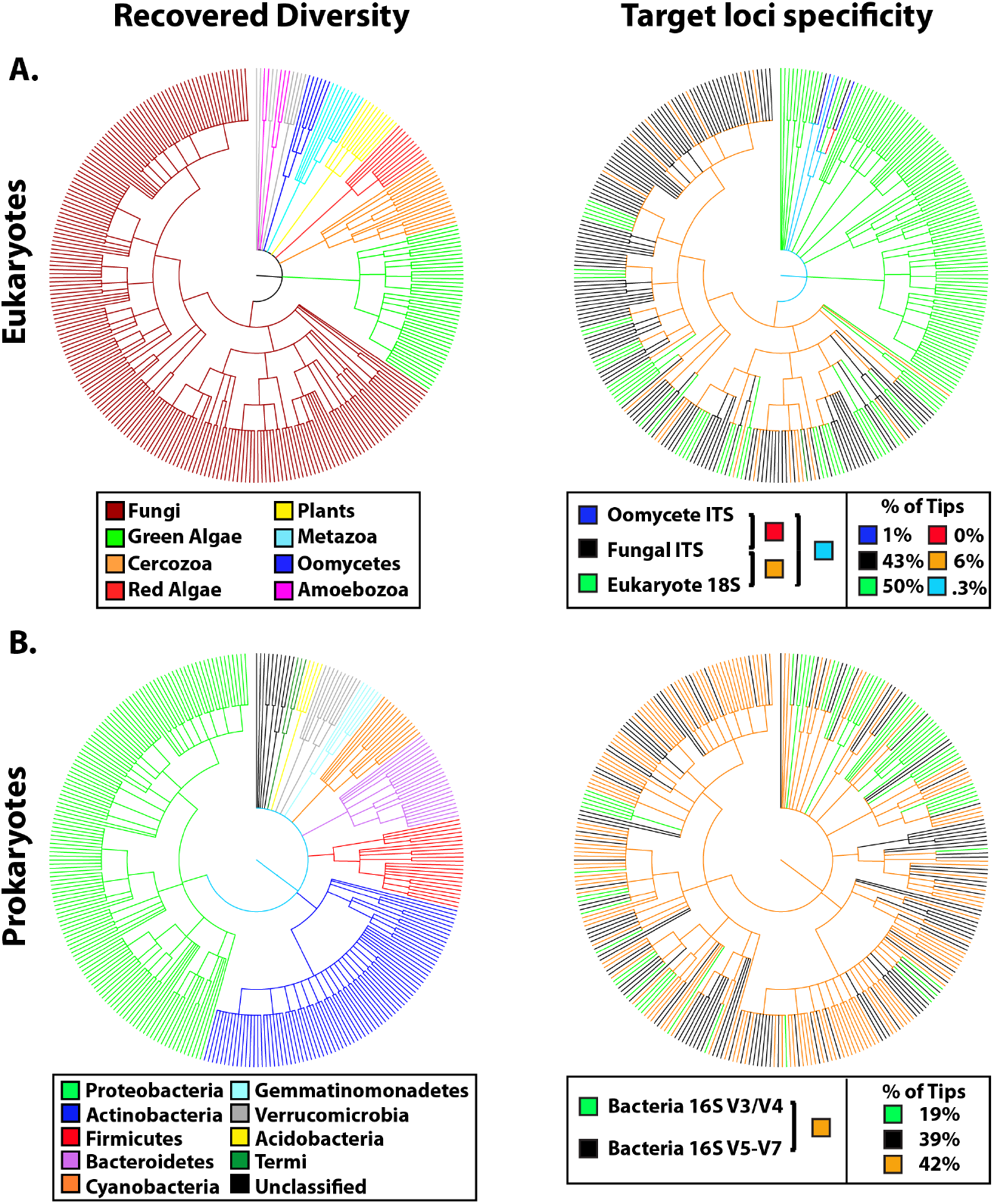
A comprehensive overview of highly diverse *A. thaliana* leaf microbiomes revealed by parallel amplicon sequencing of 8 loci targeting eukaryotic and prokaryotic microbes. Tree branches represent recovered genera and are colored by taxonomy (left, “Recovered Diversity”) and loci from which they were recovered (right, “Target loci specificity). The % of genera that were found in each dataset or by multiple datasets is presented (% of Tips). (A) Eukaryotes were targeted in 6 loci: Two regions of the 18S rRNA gene (V4-V5 and V8-V9), two regions of the fungal ITS (ITS 1 and 2) and two regions of the oomycete ITS (ITS 1 and 2). The 18S loci revealed the broadest diversity but was complemented by fungi and oomycete-specific primer sets which had more detailed resolution within these groups. (B) 2 loci targeting prokaryotes: Two regions of the 16S rRNA gene (V3-V4 and V5-V7) that amplify mostly bacteria revealed a largely overlapping diversity profile complemented by unique discovery of taxa from each of the two target regions.

### AmpStop: An “R” package for quick design of blocking oligos for any non-target organism

A key advantage of using standard oligomers as a tool to block amplification is that many design options can be tested rapidly and at low cost using standard PCR techniques. A limit on rapid implementation in labs could be the design step, where some computational know-how is required. To reduce this burden, we created “AmpStop”, an “R” package to automate design of blocking oligos. AmpStop can be used by anyone with R and BLAST+ installed on their computer. It requires as input only the amplified non-target region (for example the host ITS1 sequence) and a target sequence database that is BLAST-formatted. Three functions enable users to within minutes generate a list of all possible blocking oligos, a figure showing how many times each oligo “hits” target templates and other useful metrics of specificity (**Fig 7A–7C**) and a list of the most promising blocking oligo pairs. Since the design of peptide nucleic acid clamps follows practically identical steps^16^, the package can also be used to design them. AmpStop and detailed instructions on its use and interpretation of results is freely available on GitHub (https://github.com/magler1/AmpStop).

**Figure 7.**
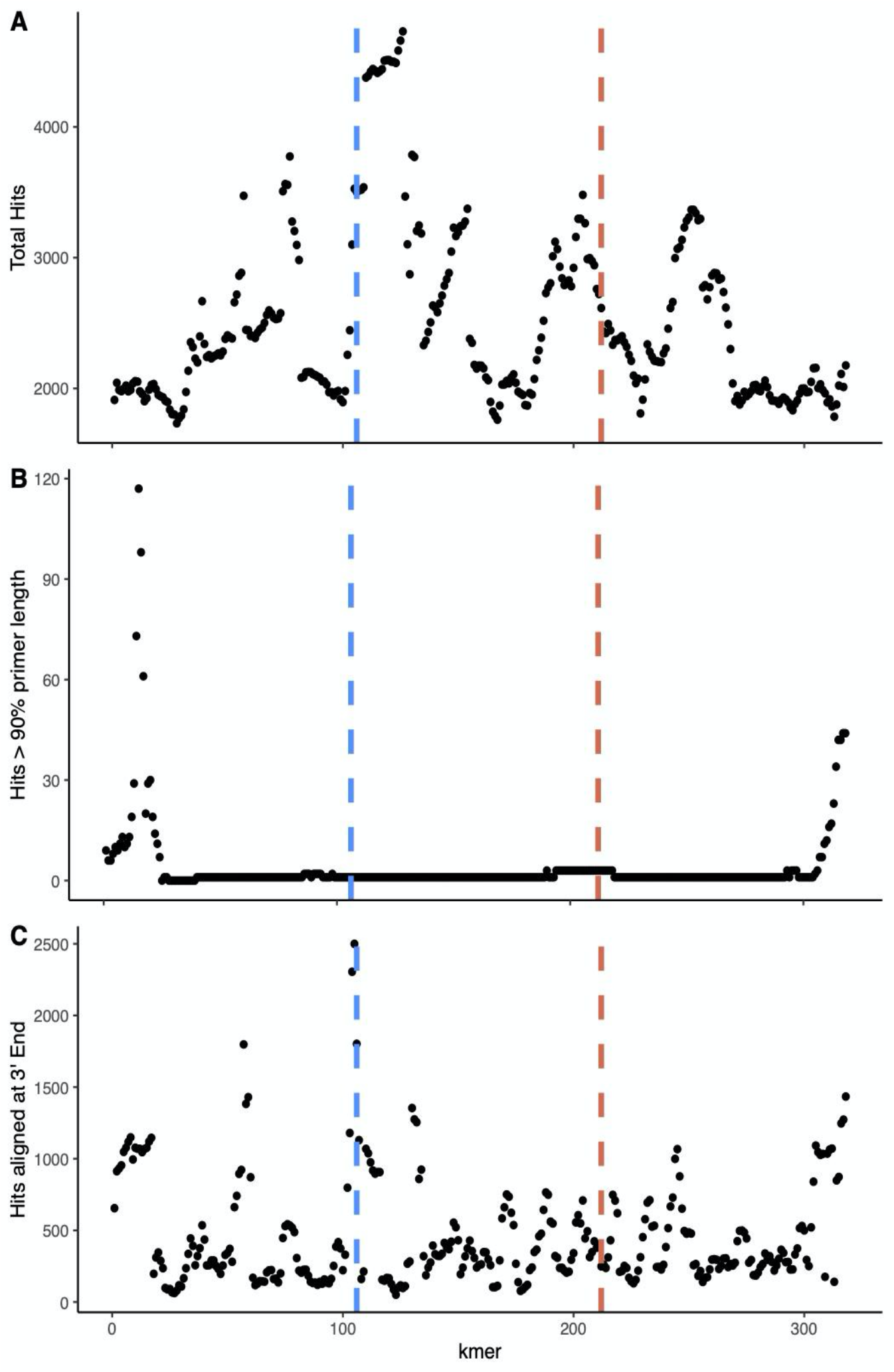
Example output from the “R” package AmpStop when given as input the *A. thaliana* ITS1 sequence (non-target) and the UNITE fungal ITS database (target). The x-axis represents all possible 30-base oligomers (candidate blocking oligos) along the length of the non-target sequence. The sequence of the oligos are produced in a separate file. **The y-axes show** three complementary measures of how likely each candidate oligo is to amplify target sequences, where a hit represents an alignment of the oligo a target. The best candidates will minimize hits and thus be highly specific to non-targets. (A) The total number of hits of each candidate to the target database. Those least likely to amplify targets will have few hits. However, not all hits are equally problematic. Thus, (B) shows oligos that hit a sequence in the target database along >90% of its length, which would increase the chance of amplifying a target organism and (C) shows hits aligned at the 3’ end, which are especially problematic because this is the site of polymerase binding. The blue and red guidelines represent 1/3 and 2/3 of the length of the non-target, respectively, and forward and reverse blocking oligos are probably best chosen before and after these lines, respectively.

## Discussion

Amplicon sequencing of phylogenetically or functionally informative loci has become an indispensable technique in a variety of biology-related fields because its targeted approach (compared to untargeted approaches like metagenomics) enables in-depth diversity characterization with accurate annotation using specialized databases^23^. It has revealed that microbial community structuring is more complex than previously thought and suggested extensive interactions between (a)biotic factors and microbes^24^ and between microbes even across kingdoms^9,11^. As we have shown here, recent advances have made sequencing up to 8 loci in parallel possible, drastically increasing throughput and diversity resolution. This will be important to add certainty to systems-scale investigations of factors contributing to microbial community structures. On the other hand, the use of “universal” primers has the disadvantage that highly abundant microorganismal or host DNA are often strongly amplified, sacrificing read depth and masking diversity^25^.

A previously described method to address this problem are peptide nucleic acid “clamps” that are highly specific to non-target templates and which physically block their amplification^16^. These clamps work efficiently even in single-step amplifications, but their production is relatively expensive, which would limit rapid development and deployment of multiple clamps for new loci, for blocking multiple non-targets and add major costs for high-throughput projects. For example, our current library preparation costs are estimated at about 2 Euros per library. PNAs, at about 4500 Euro/μmol would add 1.14 Euro or 57% per library. This would be for only host blocking and does not include costs of design and testing of new PNAs for new loci and new non-targets. Other approaches, like using oligonucleotide clamps modified with a C3 spacer^17^ are also costly and work best when they block the universal primer binding site. For many highly conserved target regions, the target and non-target binding sites are therefore too similar to design specific clamps.

Blocking oligos, which are cheap and flexible, therefore fill an important need for a tool that can be quickly designed and employed for different purposes (e.g., host or microbe blocking). Blocking can also be “dropped in” to practically any pipeline and do not bias results, so it is beneficial to include them when relative abundance of target and non-target DNA is unknown. Several different blocking techniques were previously placed under “blocking oligos” or “blocking primers” as umbrella terms^17^. However, we suggest using this term specifically for the blocking oligos we present here, as it most accurately describes their function.

An important question besides price and flexibility is whether blocking oliogs work as well as PNAs and other methods. Giancomo et al.^22^ tested blocking oligos that they designed for maize vs. other methods, including PNA clamps and discriminating primers. They recommended PNA clamps for 16S rRNA studies because they block without distorting microbiota profiles. Their maize blocking oligos did reduce plant amplification as efficiently as PNAs, but they distorted microbiota profiles in soil samples (notably not in leaf samples). Here, we tested the new universal plant 16S blocking oligos in leaves, in mock communities and in soil samples and observed no discernible effects on beta diversity. We did observe a desirable increase in alpha diversity in real leaf samples due to blocking the host and recovering more microbial reads. Thus, we recommend universal 16S and 18S plant blocking oligos, which block all plant species we tested and should also work in maize (**Fig S5**).

A downside common to blocking oligos, PNAs and other methods is that they need to be designed and tested for different non-targets, which can be cumbersome^22^. The R package “AmpStop”, which we make available here should ease the design process. AmpStop can also be used by researchers who do choose PNAs, as blocking oligos design essentially follows the same procedure^16^. The availability of universal plant blocking oligos that block amplification of most host species will further reduce the need to make new designs. Notably, we were not able to design universal blocking oligos for the ITS region because diversity between different plant species made it impossible find a universal blocking oligo set. On the other hand, we and others have observed that the host ITS is not efficiently amplified when there is significant target microbial DNA^26^. Thus, universal ITS blocking oligos are not as urgently needed as universal 16S blocking oligos.

Studies of leaves of the wild plant *A. thaliana* found that bacterial fraction of extracted DNA are typically very low but range up to about 25%^20^. We found that more blocking cycles (15 vs. 10) were necessary to efficiently block nontarget amplification in real leaf samples, but not in mock communities with 5% bacterial DNA. More cycles also lead to discovery of more alpha diversity in real leaf samples. This effect is most likely due to low bacterial loads in real leaf samples, so too few blocking cycles results in libraries that still contain relatively high levels of non-target contamination. This occurs because non-target template DNA can be carried over to the extension PCR and are amplified because the concatenated primers contain the universal primer sequence as the binding site (**Fig 1**). Thus, we recommend increasing the number of blocking cycles when following protocol A or B used here and when no prior information about bacterial loads is available. Alternatively, a linker sequence could be used as the binding site for concatenated primers in the second step^19^. This only amplifies amplicons from the first step, not left-over template DNA. Thus, blocking could be minimized to only a few cycles. Lundberg et al.^19^ did apply our ITS blocking oligos in their protocol, demonstrating that blocking oligos can be dropped into most two-step library pipelines.

Some host amplification can be advantageous because it can be used to quantitatively estimate bacterial loads^18^ and having quantitative information has can change inferred ecological relationships between plant microbiota^20^. When we only blocked chloroplast in the 16S V3/V4 region and allowed mitochondrial amplification, the fraction of bacterial reads was proportional to the bacterial DNA load in mock communities. At our target read depths in the V3-V4 region, mitochondria amplification did not overwhelm the bacterial diversity signal, but this will be locus-specific (in our hands in the V5-V7 region, mitochondrial amplification is more problematic). Generally, estimating bacterial loads from 16S rRNA data is not perfect because 16S copy numbers can vary drastically between bacterial species and plastid abundance per cell varies between eukaryotic species^27^. HamPCR^19^ is an alternative method utilizing single-copy host genes to gain accurate quantitative insights. While that approach is more precise, it does require more steps and may not be suitable for extremely high-throughput studies. Direct estimates from 16S rRNA data has the advantage of simplicity and throughput – we have designed dual-indexing primer sets to parallelize up to 500 samples (**File S1B**). Gaining approximate quantitative information here would allow users to quickly scan for plants, conditions and microbial interactions affecting bacterial load. Thus, we advise using fraction of bacterial reads in 16S data as an initial approximation to gain insight in many samples and then to design specific experiments using more precise measures.

## Conclusions

The realization of the immense complexity of biological systems – and our inability to adequately describe them - has led to many important, unresolved issues. For example, there is ongoing debate about what it means to view macroorganisms as holobionts, since symbiotic microbiota affect host health and fitness^28,29^. Unanswered questions also linger, like what causes host genotype-independent taxonomic conservation of plant root microbiomes over broad geographic distances^30^. Blocking oligos will help researchers to deeply and accurately resolve microbial community diversity when non-target contamination is problematic, addressing some of the current barriers to progress. Although other challenges remain, we expect this approach to equip researchers to make better hypotheses and to address currently intractable questions. These advances will thereby assist in increasing discovery of the important roles of microbiota.

## Methods

### Design of “blocking oligos” to avoid non-target template amplification

Blocking oligos were previously designed for the *A. thaliana* chloroplast (16S rRNA V3-V4 region) or mitochondria (16S rRNA V5-V7 region) and *A. thaliana* ITS1 and ITS2 regions (fungal and oomycete ITS)^11^. Primers specific to a known, non-target DNA template (“blocking oligos”) and nested inside the universal primer binding sites (**Fig 1**) were designed (see **File S1a and S1b** for all oligo and primer sequences used in this study). To design oligomers with high specificity, we adapted the approach used by Lundberg et al.^16^ for PNA clamps. In short, the region of interest (chloroplast/mitochondria 16S or ITS) from *A. thaliana* was divided up into “k-mer” sequences of length 30. We then used BLAST to search the kmers against a blast-formatted target database. The BLAST search used the following parameters which allow weak matches: percent identity 25, word size 7, evalue 100000. Candidate 30-mer blocking oligos were selected that received a relatively low number of hits. For this study, we have developed an R package, “AmpStop”, which automates this part of the process and suggests good candidates, and which is freely available on GitHub (https://github.com/magler1/AmpStop). We then selected candidates which had a high Tm (well above the universal primer binding temperatures) and which had low potential to form self-dimers or hairpins. Candidate oligomers were tested in single-step amplification of target and non-target templates for non-target specificity. The selected blocking oligomers (**File S1**) were always used in the first amplification step of library preparation (blocking cycles), resulting in shortened amplicons that could not be elongated with Illumina adapters in the second amplification step (**Fig 1B**). All databases will be made publicly available prior to publication on Figshare.

### Design of 18S blocking oligos for host and microbial non-targets

To reduce both *A. thaliana* and *A. laibachii* amplification in the 18S regions we designed additional blocking oligos for both of these organisms (**File S1**). We tested them by preparing 18S amplicon libraries from two mock communities consisting of *A. thaliana* (97% or 87%), *A. laibachii* (0 or 10%)*, Sphingomonas* sp. (1.5%), *Bacillus* sp. (1.5%) and 0.001% to 1% of target *Saccharomyces* cerevisiae. (**Table S2**). We then used primers targeting the *Saccharomyces* sp. 18S (V4 Fwd/Rev: AACCTTGAGTCCTTGTG/AATACGCCTGCTTTG V9 Fwd/Rev: GTGATGCCCTTAGACG/ACAAGATTACCAAGACCTC) with qPCR to relatively quantify target *S. cerevisiae* in the libraries generated with and without blocking oligos (**Fig 4B**).

### Design of universal plant blocking oligos for 16S and 18S rRNA loci

To design blocking oligos that could be used for multiple plant species we used the same approach as described above. In short, we used chloroplast sequences from multiple plant species that spanned the phylogentic tree of plants (**see Table S4**) as input for the AmpStop package and checked where the results overlapped between species. The resulting blocking oligos were tested in a one-step PCR protocol for their specificity against genomic DNA from various plant species and bacterial mixes (**Fig S5**). The blocking oligo pair BloO_16S_F5 and BloO_16S_R1 was chosen for further analysis, since it hit most of the plant species tested but at the same time amplified none of the bacterial mixes. These selected oligomers were used in the blocking cycles for library preparation from multiple plant species.

### Testing A. thaliana blocking oligos against mock communities

We tested blocking oligos designed to block *A. thaliana* in two loci from each of bacteria (16S rRNA V3-V4 and V5-V7), fungi (ITS1 and 2) and oomycetes (ITS1 and 2) using mixed kingdom mock microbiomes. The simulated host-associated microbiomes consisted of 5% of a mix of the mock microbiomes and 95% *A. thaliana* genomic DNA. For each template sample, 6 separate PCR reactions were prepared, one targeting each locus. We also tested the effect of variations on the amplicon sequencing library preparation method. We compared PCR performed in one step (35 cycles, no blocking) or two steps (10 then 25 cycles or 25 then 10 cycles). For two-step preparations, the primers used in the first step consisted of unmodified universal amplification primers (**Fig. S1**). For single-step preparations and for the second step in two-step preparations, primers were a concatenation of the Illumina adapter P5 (forward) or P7 (reverse), an index sequence (reverse only), a linker region, and the universal primer for the region being amplified (**Fig 1** and **Fig S1**). Sequences and details of all primers used can be found in **File S1a** and details on PCR, library preparation and sequencing, as well as the steps to generate OTU tables and taxonomy from raw multi-locus data can be found in the **Supporting Methods (Protocol A)**. We summarized bacterial, fungal and oomycete OTU tables by taxonomic ranks, converted abundances to relative values and plotted the genus- and order-level taxonomic distribution directly from this data with the package ggplots2 in R. To analyze the percent reduction in host plant-associated reads when blocking oligos were employed, we considered the relative abundance of reads associated with the class “Chloroplast” or the order “*Rickettsiales”* in the 16S OTU tables and reads in the kingdom *“Viridiplantae”* in the ITS OTU tables in samples with *A. thaliana* DNA and with and without blocking oligos. We also checked whether the non-indexed step of the 2-step library preparation approach results in sample cross-contamination by sequencing three negative control libraries (two blank samples carried through DNA extraction and one PCR water control) by adding the whole volume of the libraries to the combined sequencing pool. These negative controls were prepared in parallel with 381 other plant samples (**Supporting Notes**).

### Testing “universal” plant blocking oligos in natural leaves and mock communities

16S and 18S universal plant blocking oligos were tested using five leaves from five plant species collected from different experiments. The plant species represent five plant orders spanning monocots and dicots (see **Table S5**). All plant leaves were naturally grown outside without artificial addition of any microorganism. Details on the DNA extraction can be found in the **Supplementary Methods**. The 16S universal blocking oligos were additionally tested for how they affect bacterial diversity distribution against a mixed microbial community standard (ZymoBIOMICS microbial community DNA standard) and against three different soil DNA extracts. For testing if bacterial load information is maintained, we quantified concentrations of axenic platn DNA extracts and combined it with the Zymo standard to create genomic DNA mixes (0%, 0.01%, 0.05%, 0.1%, 0.5%, 1%, 15% and 25% microbial genomic DNA).

Libraries were prepared with either 10 or 15 blocking cycles and 25 or 20 extension cycles, respectively. In short, the templates were amplified in the blocking cycles including the universal 16S primers as well as the blocking oligos. The product of this first PCR was then purified and amplified in the extension cycles using concatenated primers. The extension step used concatenated primers similar to before but both primers had unique index sequences. Sequences and details of all primers used can be found in **File S1b** and details on PCR, library preparation and sequencing, as well as the steps to generate ASV tables and taxonomy from raw data can be found in the **Supporting Methods (Protocol B)**.

### *8-locus amplicon sequencing with blocking oligos to fully characterize* A. thaliana *leaf microbiome diversity*

We next expanded the multi-locus approach to more completely cover eukaryotic microbial diversity by including two additional 18S rRNA gene loci (V4-V5 and V8-V9, **Fig. 1a** and **Table S1** - primer sequences are available in **File S1**). With the expanded target set, we characterized the phyllosphere microbiome of *A. thaliana* leaves infected with the oomycete pathogen *Albugo laibachii*. Whole leaves (defined as a single whole rosette) or endophytic fractions of leaves (defined as in^11^) were collected in the wild (a total of 18 samples - 9 whole leaf, 9 endophyte) and were immediately frozen on dry ice. DNA extraction was performed as described previously^11^. Library preparation, sequencing and analysis was performed as in the optimized protocol with blocking oligos. To provide a complete and concise picture of the diversity of microbiota inhabiting *A. thaliana*, we combined the data from all samples. To visualize data, we assigned taxonomy to OTUs and generated two phylogenetic trees where branches represent unique genera. Trees were generated based on the taxonomic lineages (*not* phylogenetic relatedness of OTUs or genera) with the ape package in R and output as newick files^31^. The trees were uploaded to iTOl v3.1^32^ to color branches by taxonomy or by targeted regions. The first tree (**Fig. 3A**), for Eukaryotes, includes data from the 18S rRNA and ITS targeted regions. The second tree includes data from the 16S rRNA targeted regions.

## Declarations

### Ethics approval and consent to participate

Not applicable

### Consent for publication

Not applicable

### Availability of Data and Material

Universal primer sequences, sequencing primers, blocking oligos and concatenated primer sequences are all provided in File S1a and S1b.

Scripts used to generate ASV tables from the raw data, as well as OTU/ASV tables and metadata files to recreate the main figures are being made publicly via Figshare: https://figshare.comIprojectslObtaining_deeper_insights_into_microbiome_diversity_using_a_simple_method_to_block_host_and_non-targets_in_amplicon_sequencing_/89504 Raw sequencing data is publicly available as a NCBI projects (*PRJNA420016 and PRJNA663775*).

AmpStop is freely available on GitHub (https://github.com/magler1/AmpStop).

### Competing Interests

The authors declare that they have no competing interests.

### Funding

TM and MTA are supported by the Carl Zeiss Stiftung *via* the Jena School for Microbial Communication. MMR is supported by the International Leibniz Research School. EK, JA, SH and ND were supported financially by the Max-Planck Gesellschaft and the University of Tuebingen. AM Acknowledges the financial support from the European Research Council (ERC) under the DeCoCt research program (grant agreement: ERC-2018-COG 820124). SH and ND were supported financially by the Max-Planck Gesellschaft.

## Acknowledgements

We wish to thank Ariane Kemen and Jonas Ruhe for providing *P. capsici* and *Pseudozyma* sp. isolates and Marie Harpke and Prof. Erika Kothe for providing soil DNA. We also thank the MPIPZ genome center for implementing our custom sequencing protocol and Carl-Eric Wegner, Stefan Riedel and Prof. Kerstin Küsel for making their sequencing equipment and knowledge available to us.

## Notes

https://figshare.com/projects/Obtaining_deeper_insights_into_microbiome_diversity_using_a_simple_method_to_block_host_and_non-targets_in_amplicon_sequencing_/89504

https://github.com/magler1/AmpStop

### Competing Interest Statement

The authors have declared no competing interest.

