## Supporting Information for "Obtaining deeper insights into microbiome diversity using a simple method to block host and non-targets in amplicon sequencing"

**Figure S1- S12**
**Table S1-S7**
**Supporting Notes**

### Supporting Methods

### Supporting Figures

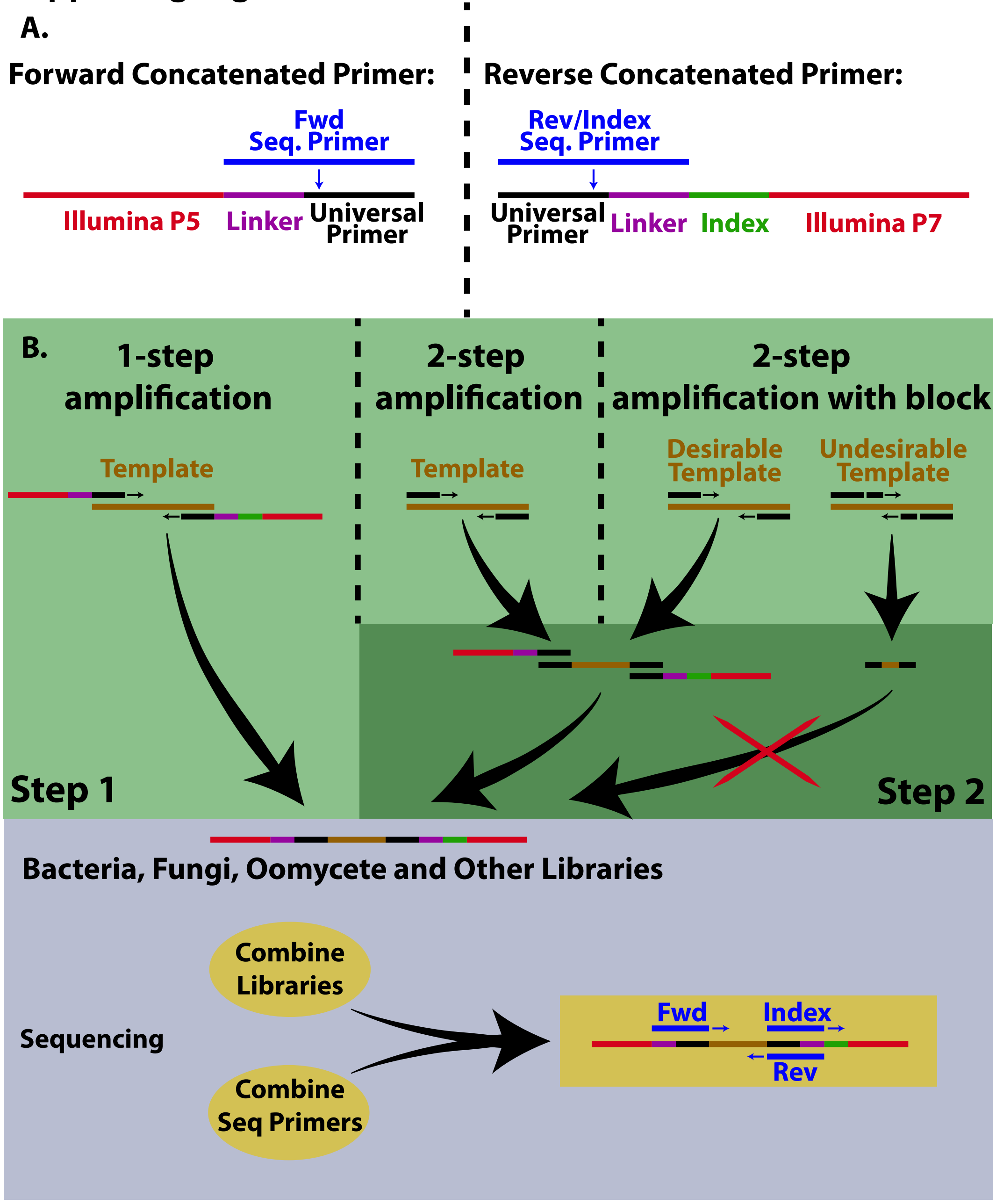


**Figure S1. Amplification and sequencing strategies compared in this study.** A. The forward and reverse concatenated primers include Illumina adapter, an index to identify which samples the library derived from (reverse only), a linker region and the universal primer. Three sequencing primers are complementary to the linker/universal primer region. B. Three amplification and sequencing strategies were compared: 1-step amplification with concatenated primers, 2-step amplification with universal primers followed by a second step with the concatenated primers, and 2-step amplification including blocking oligomers to reduce amplification of undesirable template, such as non-microbial host DNA. C. Sequencing was carried out by combining libraries from all samples and groups of microorganisms that were amplified and sequencing in three steps with a mix of sequencing primers. One set of sequencing primers are required for each universal primer set that is used in library preparation.

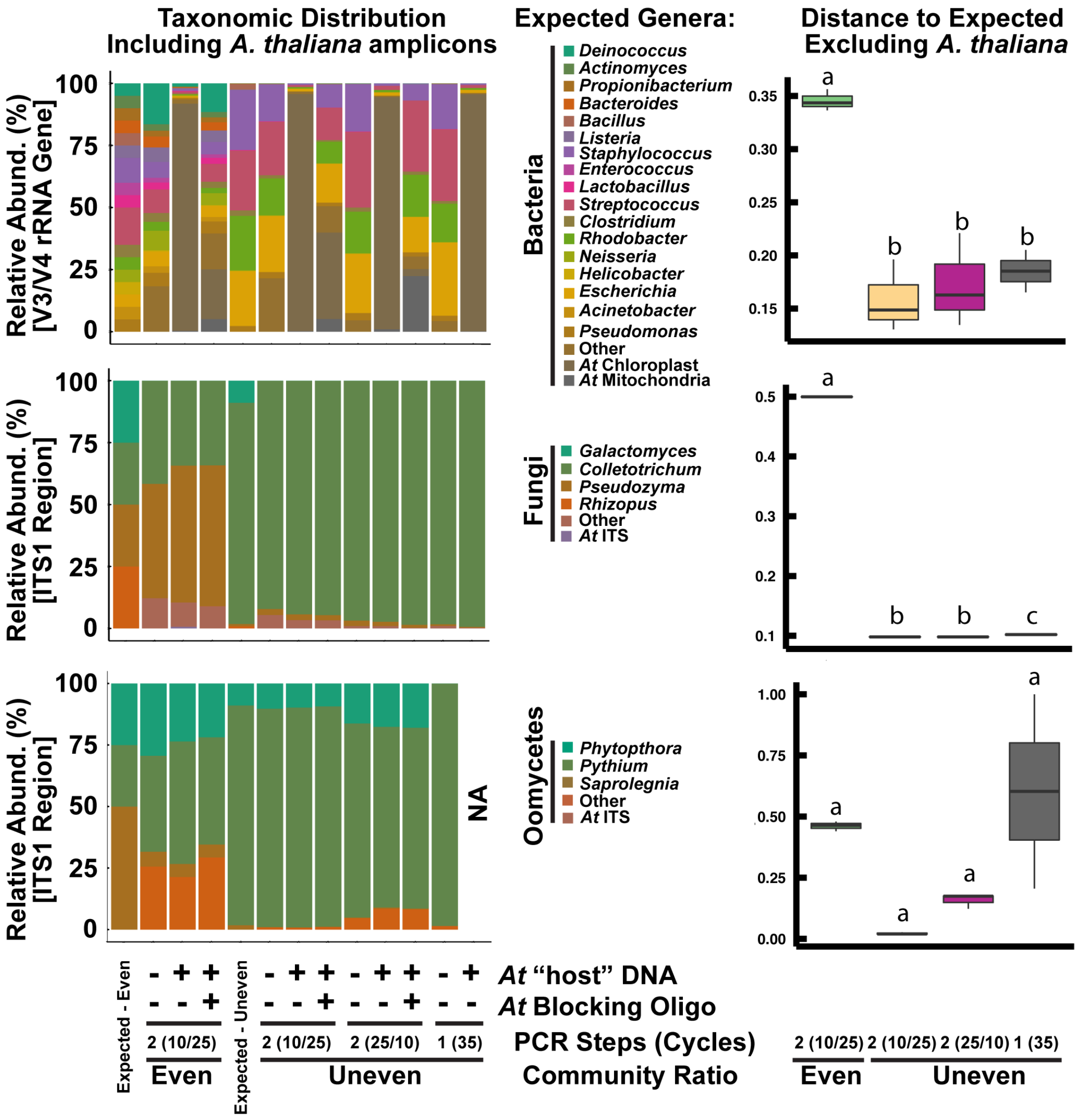


**Figure S2. Reproducible and accurate characterization of mock communities of bacteria, fungi, and oomycetes by amplicon sequencing (complementary loci to Figure S4).** A. Observed taxa at the genus level in sequenced mock communities closely matched expected communities. The taxa "Other" is primarily non-target amplification from *A. thaliana* "host" DNA that was added to test blocking oligomers which prevent "host" DNA amplification. "NA" indicates a sample where sequencing depth was too low after subsampling to be included. B. Distance (Bray-Curtis distance based on relative abundance of genus-level taxa) of sequenced communities from the expected distribution where 0 is identical and 1 is unrelated. Even or staggered revers to the distribution of the organisms in the mock communities (see expected distributions in A). PCR Steps refers to a 1-step (35 cycles with concatenated primers) or 2-step (10 or 25 cycles with standard primers followed by 25 or 10 cycles with extension primers containing Illumina adapters) amplification protocol. Letters indicate $p < 0.05$ (FDR-corrected) based on pairwise t-tests between groups.

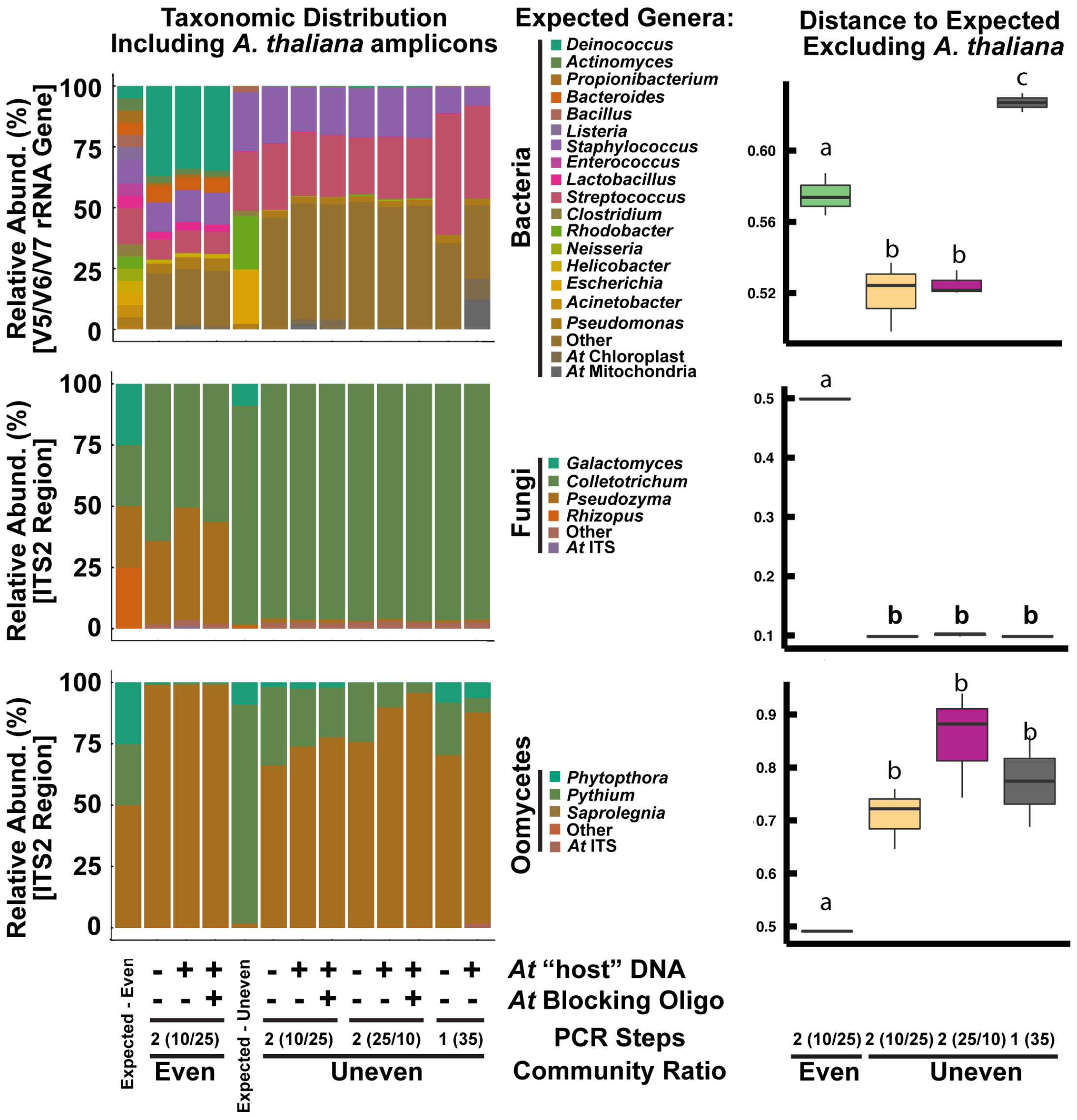


**Figure S3**. **Reproducible and accurate characterization of mock communities of bacteria, fungi, and oomycetes by amplicon sequencing (complementary loci to Figure S3).** A. Observed taxa at the genus level in sequenced mock communities closely matched expected communities. The taxa "Other" is primarily non-target amplification from *A. thaliana* "host" DNA that was added to test blocking oligomers which prevent "host" DNA amplification. "NA" indicates a sample where sequencing depth was too low after subsampling to be included. B. Distance (Bray-Curtis distance based on relative abundance of genus-level taxa) of sequenced communities from the expected distribution where 0 is identical and 1 is unrelated. Even or staggered refers to the distribution of the organisms in the mock communities (see expected distributions in A). PCR Steps refers to a 1-step (35 cycles with concatenated primers) or 2-step (10 or 25 cycles with standard primers followed by 25 or 10 cycles with extension primers containing Illumina adapters) amplification protocol. Letters indicate $p < 0.05$ (FDR-corrected) based on pairwise t-tests between groups.

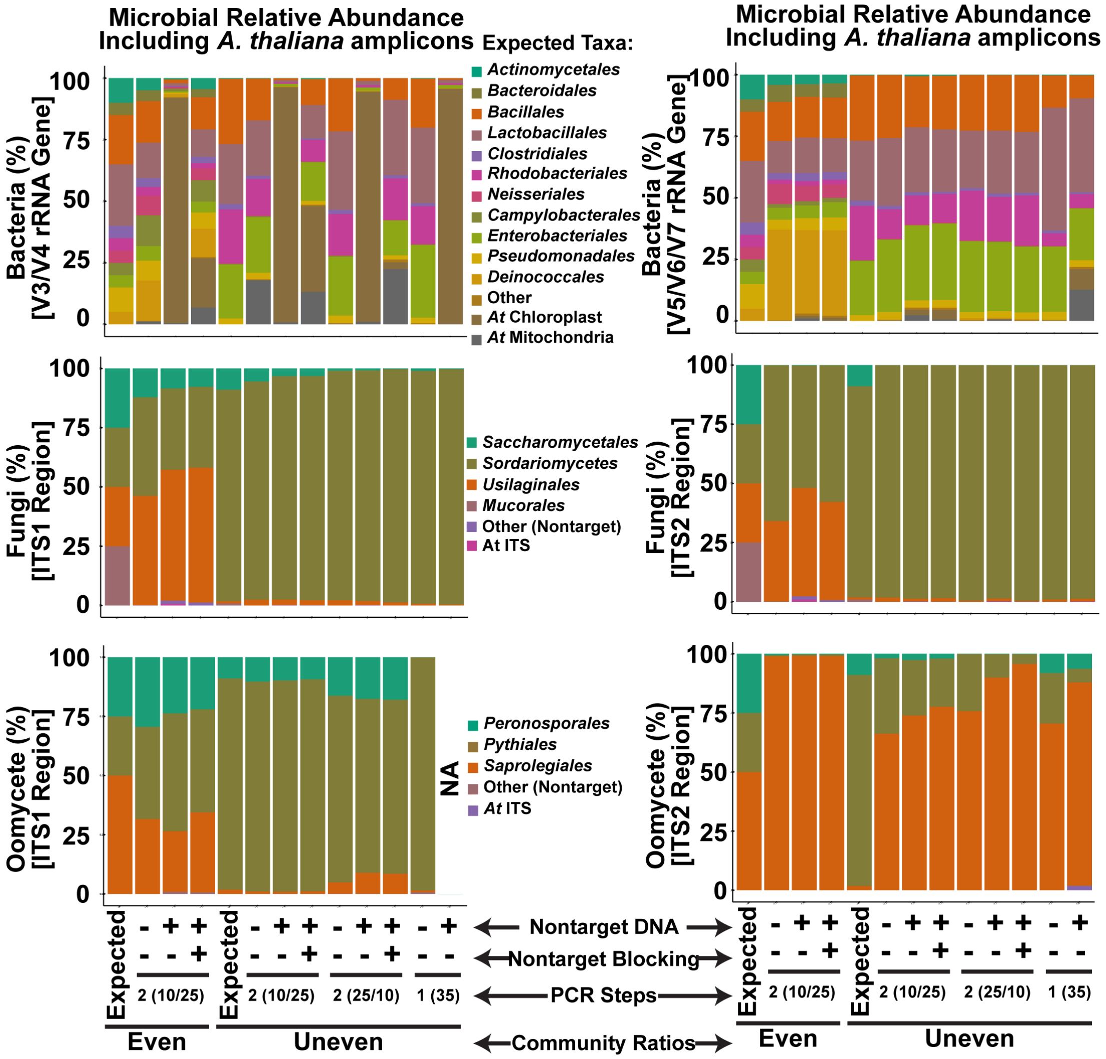


**Figure S4. Reproducible and accurate characterization of mock communities of bacteria, fungi, and oomycetes by amplicon sequencing (order-level taxonomy).** Observed taxa at the order level in mock communities sequenced at 6 loci closely matched expected communities. The taxa "Other" is primarily non-target amplification from *A. thaliana* "host" DNA that was added to test blocking oligomers which prevent "host" DNA amplification. "NA" indicates a sample where sequencing depth was too low after subsampling to be included.

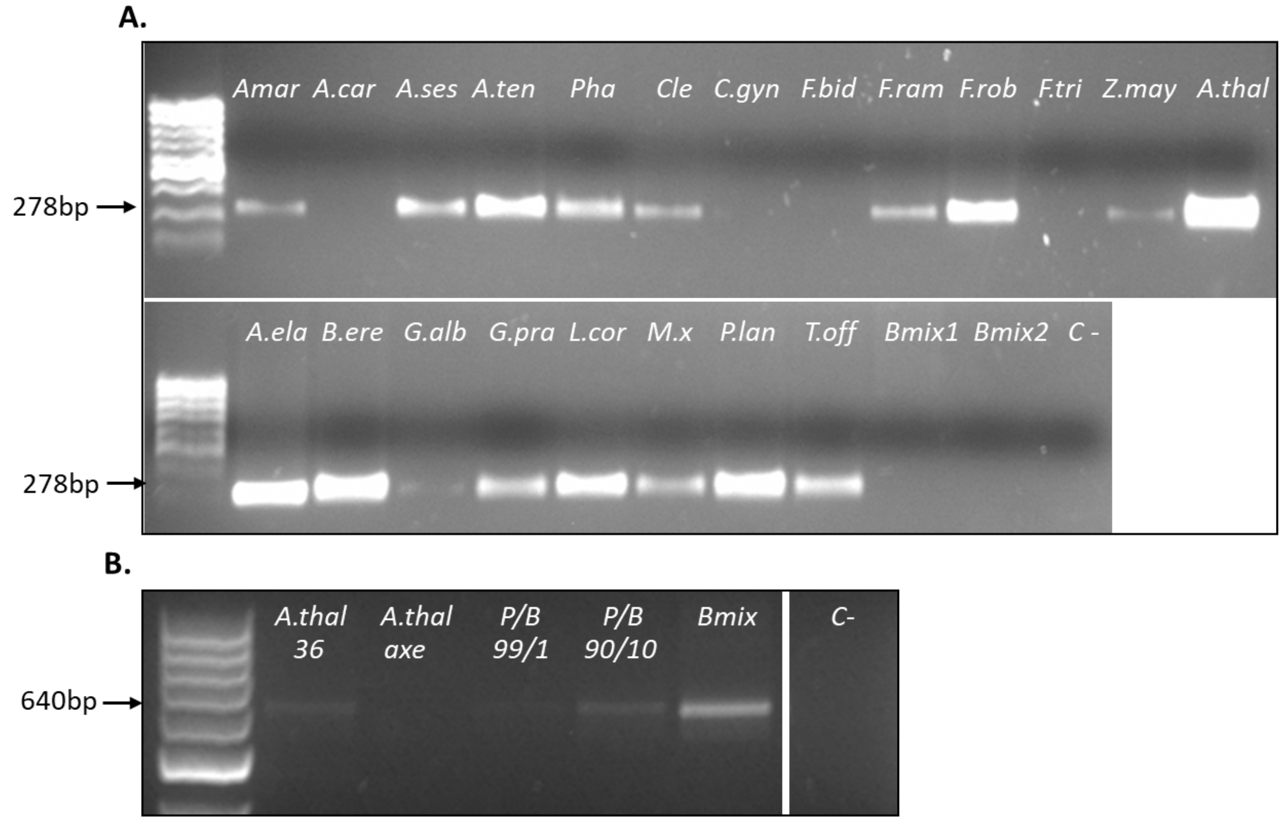


**Figure S5: A. Blocking oligos amplified most of the tested plant species but not the bacterial mixes in a single step PCR.** A wide variety of different plant species were used to test initially if the Blocking Oligos F5/R1 successfully amplify plant chloroplast DNA and at the same time do not amplify bacterial DNA. Full names and heritage of the plants used can be found in table SI4. **B. In the final two-step PCR protocol B the use of the blocking oligos F5 and R1 results in one lane in a plant sample with bacteria and in no lane in samples without bacteria.** Before the eventual sequencing run the protocol was tested extensively resulting in the desired gel picture in the end: In the plant with bacteria there is only one lane (the bacterial one), in the axenic plant there is no lane (because the amplification of the plant DNA is blocked by the blocking oligos) and the sample with only bacteria (positive control) also shows a clear lane.

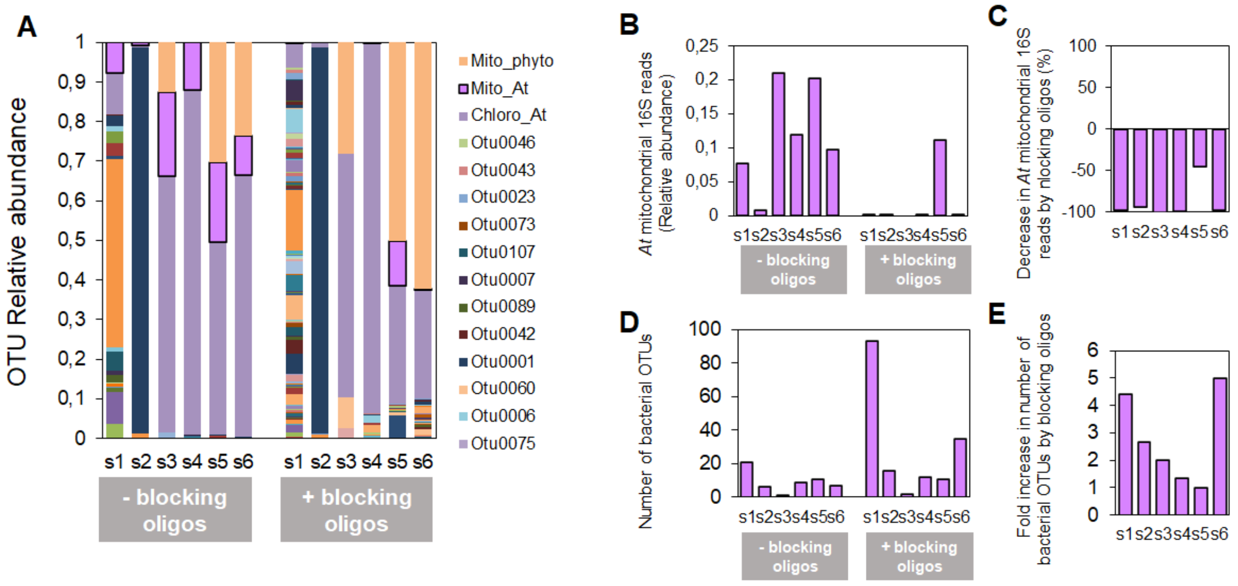


**Figure S6. Partially blocked amplification of *Arabidopsis thaliana* (*At*) mitochondrial 16S by sequence specific blocking oligos during bacterial microbiome analysis improves diversity detection.** Sequencing of the bacterial 16S rRNA v5-v7 region was conducted on six *At* plants collected from natural populations (samples s1 to s6). The effect of PCR blocking oligos designed to specifically block *At* mitochondrial 16S amplification was tested by adding (+ blocking oligos) or not (- blocking oligos) the blocking oligos in the first PCR step of the procedure. (A) Relative abundance of bacterial taxa (OTUs) and organellar 16S reads. “Mito_phyto”: Mitochondrial 16S sequences from Phytophtora; “Mito_At”: Mitochondrial 16S sequences from *At*; “Chloro_At”: Chloroplastic 16S sequences from *At*. (B) Relative abundance of *At* 18S reads with/without PCR blocking blocking oligos. (C) Percentage of decrease in the relative abundance of *At* mitochondrial 16S reads upon addition of PCR blocking blocking oligos. (D) Number of bacterial taxa (OTUs) detected in the samples with and without mitochondrial 16S PCR blocking blocking oligos. (E) Fold increase in the number of bacterial taxa (OTUs) detected in the samples upon addition of mitochondrial 16S PCR blocking blocking oligos.

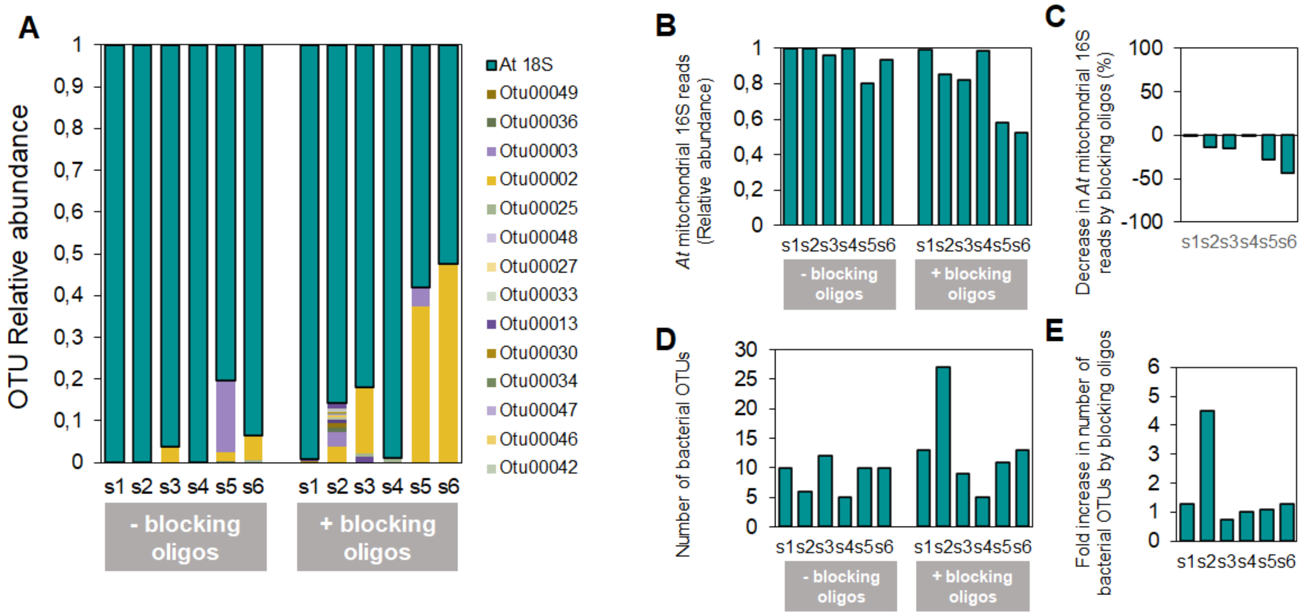


**Figure S7. Partially blocked amplification of *Arabidopsis thaliana* 18S rRNA by sequence specific blocking oligos during micro-eukaryotic microbiome analysis improves diversity detection.** Sequencing of the eukaryotic 18S rRNA v9 region was conducted on six *At* plants collected from natural populations (samples s1 to s6). The effect of PCR blocking oligos designed to specifically block *At* 18S amplification was tested by adding (+ blocking oligos) or not (- blocking oligos) the blocking oligos in the first PCR step of the procedure. (A) Relative abundance of micro-eukaryotic taxa (OTUs) and plant 18S reads. “At_18S”: 16S rRNA sequences from *At.* (B) Relative abundance of *At* 18S reads with/without PCR blocking blocking oligos. (C) Percentage of decrease in the relative abundance of *At* 18S reads upon addition of PCR blocking blocking oligos. (D) Number of micro-eukaryotic taxa (OTUs) detected in the samples with and without *At* 18S PCR blocking blocking oligos. (E) Fold increase in the number of micro-eukaryotic taxa (OTUs) detected in the samples upon addition of *At* 18S PCR blocking blocking oligos. Histograms bars depict values for individual samples.

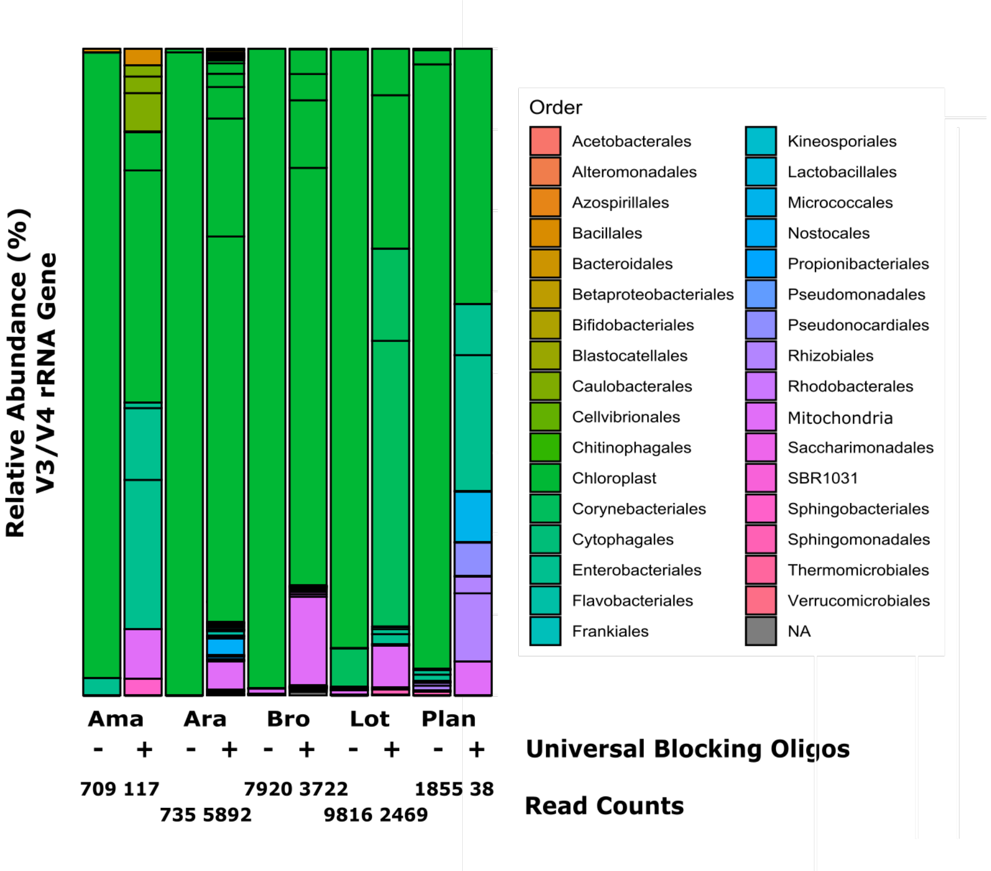


**Figure S8. Characterization of microbial communities on "real" plant samples already showing that the use of blocking oligos for 15 cycles increases the amount of bacterial reads retrieved.** For all of the five test species the use blocking oligos leads to a higher fraction of bacterial reads recovered from those samples. For the samples from *Amaranth spec.* (Ama) and *Plantago lanceolata* (Plan) the overall number of reads was low and they were excluded from further analysis.

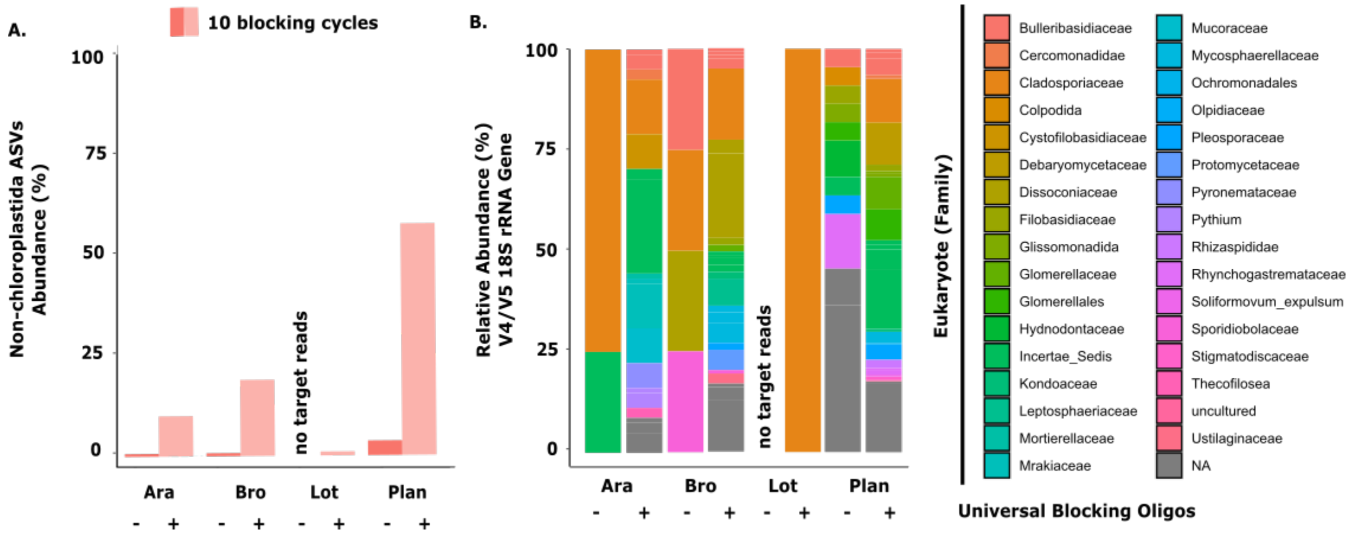


Figure S9: **Universal 18S blocking oligos successfully block for recovered eukaryotic species and therefor increasing sequencing depth.** (A) Percentage of reads assigned to ASVs other than chloroplastida (non-chloroplastida ASVs) with 10 blocking cycles. The use of blocking oligos leads to higher recovery of eukaryotic ASVs. (B) Taxonomic distribution in samples of different plant species with and without blocking oligos (10 cycles).

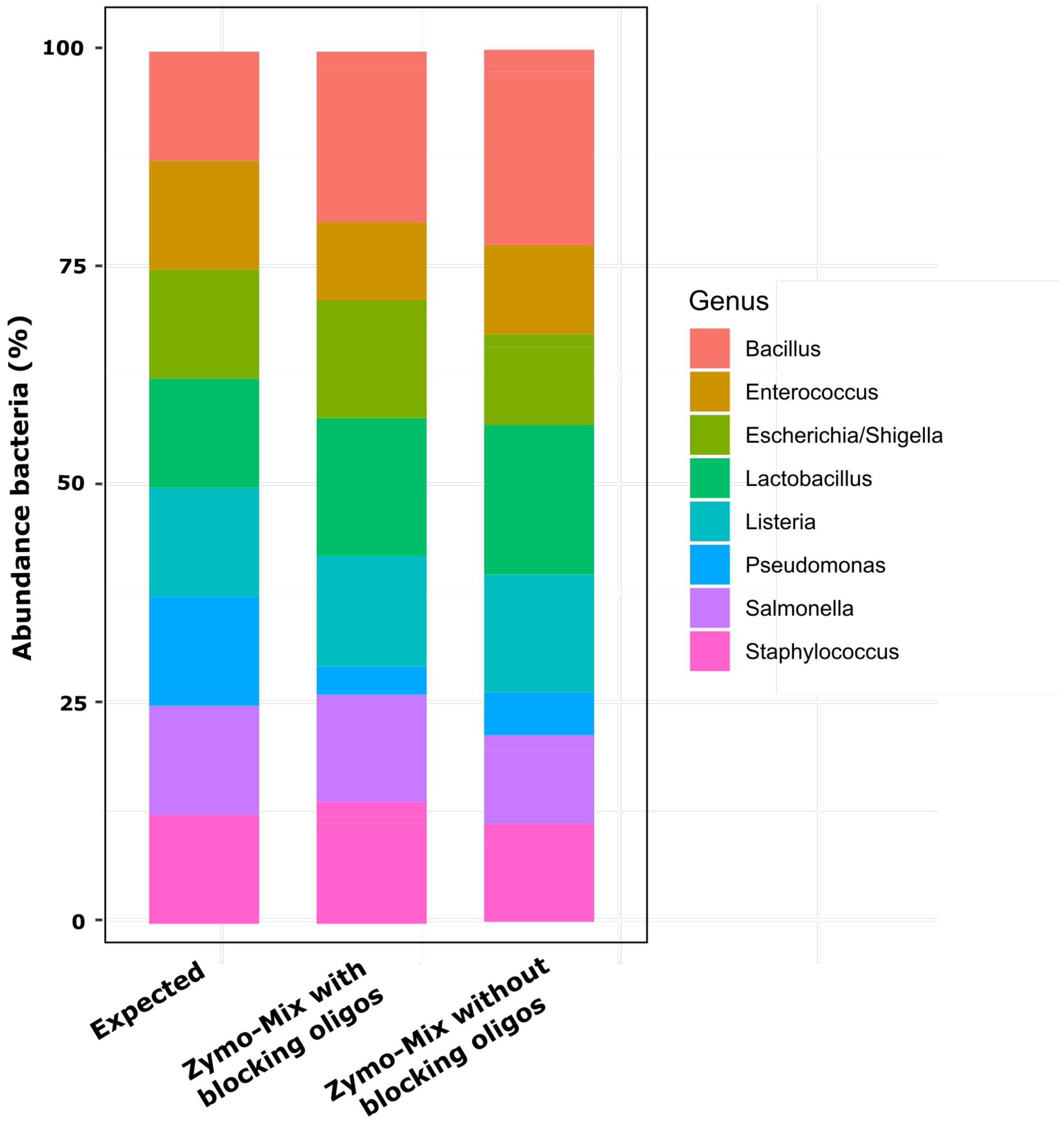


**Figure S10. Zymo control barchart: Use of universal blocking oligos does not obscure the microbial diversity recovered.** Two control samples were added in the sequencing run testing the universal blocking oligos, which including 1uL of the Zymo Microbial mix, universal bacterial primers and either universal 16S blocking oligos or not. Based on the information of Zymo we created “Expected”-barchart, which is the barchart we would expect if the sequencing was 100% accurate. As seen in the Figure, the microbial diversity in general shifts a bit in favour of *Bacillus* and unfavour of *Pseudomonas*, but this effect is similar regardless of the use of blocking oligos or not.

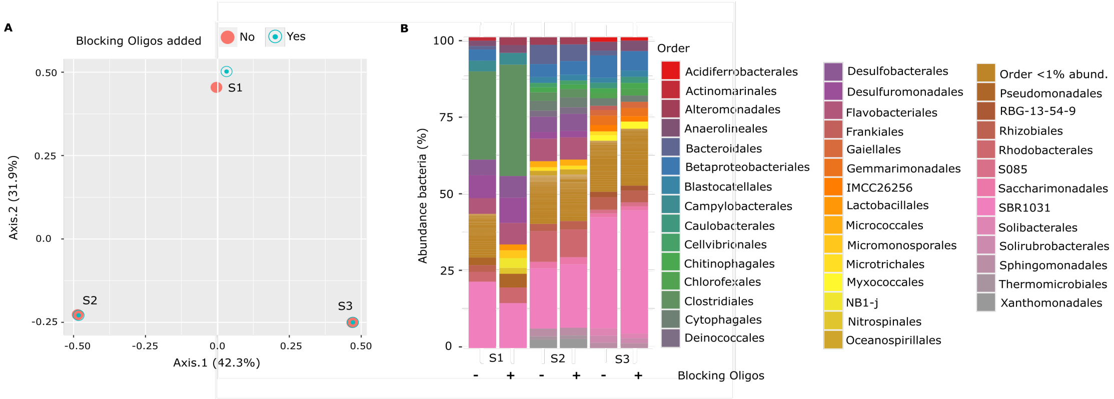


**Figure S11. Universal blocking oligos added to soil samples (DNA extracted from soil) do not significantly change the bacterial composition of the samples.** Three soil samples were added in the sequencing run. All three were sequenced either with universal blocking oligos or without. The microbial composition was analysed and plotted as MDS (distance "bray") (A) as well as barcharts showing the sequenced bacteria on order level that are more abundant than 1% (B). Both figures show that the bacterial composition change only marginally within the samples amplified with or without blocking oligos added to the first PCR.

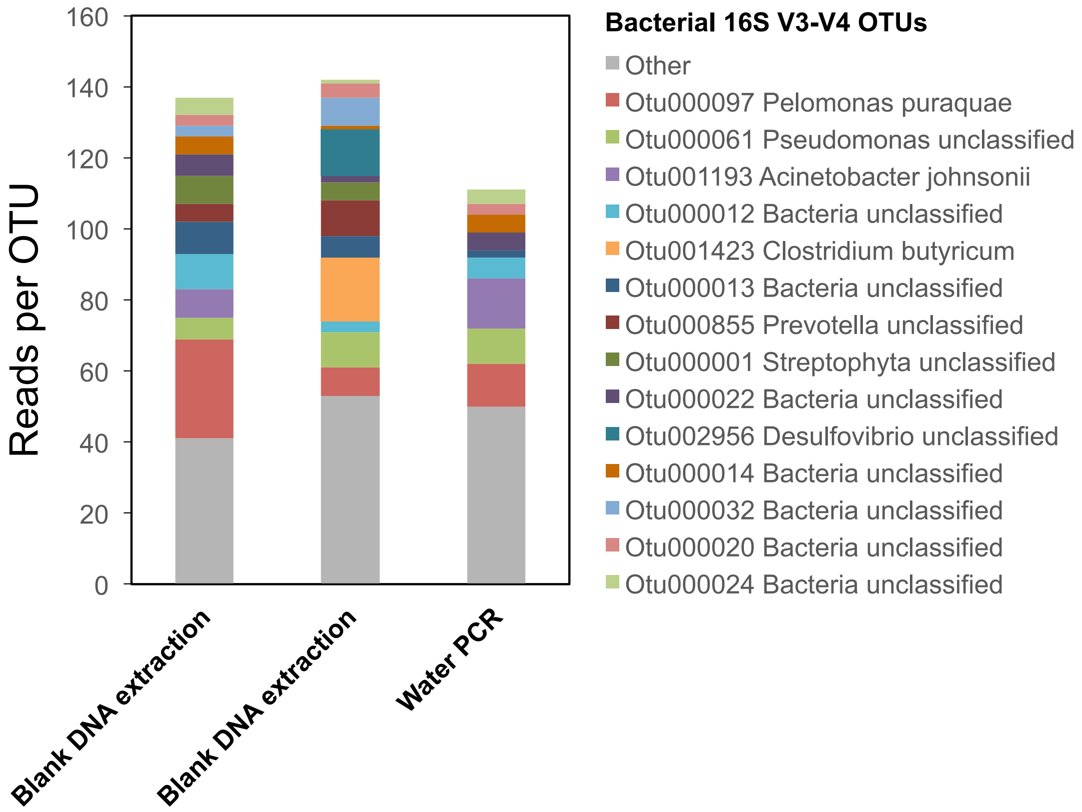


**Figure S12. Low numbers of bacterial 16S V3-V4 reads detected in negative controls indicate a low level of environment-derived contamination and a low level of cross contamination between samples.** Three negative controls including samples from two blank DNA extractions ("Blank DNA extraction") and one negative PCR ("Water PCR") were included in a sequencing run of 381 samples from *A. thaliana* leaves. The samples were treated and analyzed as described previously. The number of bacterial 16S V3-V4 sequences per OTU is shown, OTU "Other" indicates reads belonging to other low abundance OTUs (< 10 total reads)..

### Supporting Tables

**Table S1.** Gene regions and universal primers utilized in this study.

| Kingdom | Locus | Universal Primers | References |
|---|---|---|---|
| Bacteria | 16S rRNA V3/V4 | B341F/B806R | [Muyzer, Applied and Environomental Microbiology, 1993] / [Caporaso et al., PNAS, 2010] |
| | 16S rRNA V5/V6/V7 | B799F/B1194R | [Chelius and Triplett, Microbial Ecology, 2001] / [Bodenhausen et al., PlosONE, 2013] |
| Fungi | ITS1 | ITS1F/ITS2 | [Gardes and Bruns, Mol Ecology, 1993] / [White et al, PCR Protocols book chapter, 1990] |
| | ITS2 | fITS7/ITS4 | [Ihrmark et al., Micorbiology Ecology, 2012] / [White et al, PCR Protocols book chapter, 1990] |
| Oomycetes | ITS1 | ITS1O/5.8s-O-R | [Thines, J. Phytopathology, 2006] / (designed for this study, supplementary note) |
| | ITS2 | 5.8s-O-F/ITS4 | (designed for this study, supplementary note) / [White et al, PCR Protocols book chapter, 1990] |
| Other Eukaryotes | 18S rRNA V4/V5<br>18S rRNA V8/V9 | F566/R1200<br>F1422/R1797 | |

**Table S2.** Mock communities used to test blocking oligos in the 18S rRNA target regions.

| Component | A | B |
|---|---|---|
| *A. thaliana* | 97% | 87% |
| *Sphingomonas* sp. | 1.5% | 1.5% |
| *Bacillus* sp. | 1.5% | 1.5% |
| *A. laibachii* | 0% | 10% |
| *Saccharomyces* sp. | 1% [**A1**] | 1% [**B1**] |
| | 0.1% [**A2**] | 0.1% [**B2**] |
| | 0.01% [**A3**] | 0.01% [**B3**] |
| | 0.001% [**A4**] | 0.001% [**B4**] |

**Table S3.** Sequences used for designing universal plant blocking oligos

| Region | Organism | Database ID | Database |
|---|---|---|---|
| 18S V4/V5 | *A. thaliana* | AC004708.14310.16107<br>JSAD01000227.3546.5328 | Silva 18S |
| | *Bromus tectorum* | GEFS01004938.10.1399 | |
| | *Amaranthus tuberculatus* | ACQK01000002.994.2789 | |
| | *Geranium* sp. | U42541.1.1729 | |
| | *Lotus japonicus* | AP004494.26295.28091<br>BABK02040944.41558.43052 | |
| | *Medicago trunculata* | MWMB01000079.55419.57036<br>AC136838.69231.70825 | |
| | *Phaseolus vulgaris* | LPQZ01042225.748.2544<br>ANNZ01004572.5334.6969 | |
| | *Plantago lanceolata* | AJ236046.1.1766 | |
| | *Brachypodium distachyon* | ADDN02000415.208502.210251 | |
| | *Zea mays* | MTTA01000006.25673421.25675221<br>AC217805.154821.156622 | |
| 16S V3/V4 | *Taraxacum officinale* | *NC_030772.1:99536-101026* | NCBI |
| | *Gallium mollugo* | NC_036970 | |
| | *Geranium incanum* | KT760575.71428.72920 | Silva 16S |
| | *Geranium phaeum* | KT760577.136627.138133 | |
| | *Geranium robertianum* | HAGH01000827.5518.7007 | |
| | *Geranium pyrenaicum* | HAGG01001037.200.1694 | |
| | *Lotus japonicus* | AP007658.33144.34619<br>BABK02041417.620402.621870 | |
| | *Lotus corniculatus* | GACB01007670.5.1415 | |
| | *Plantago maritima* | KR297244.103587.105077 | |
| | *Medicago trunculata* | AC093544.25901.27371<br>AC174362.52078.53549<br>MWMB01000001.21918288.21919749<br>MWMB01000582.3522.4980<br>MWMB01000004.2546045.2547315<br>APNO01004332.21784.23254<br>APNO01003868.437917.439230 | |
| | *Bromus tectorum* | GEFS01004945.10.1322<br>GEFS01004938.10.1399<br>GEFS01004943.10.1226 | |
| | *Phaseolus vulgaris* | DQ323045.3862.5314 | |
| | *Zea mays* | AC204691.103667.105015<br>AC209758.219396.220865 | |
| | *Arabidopsis thaliana* | JSAD01000349.12836.14249<br>JSAD01000356.15043.16472 | |

**Table S4.** Plants used for testing universal blocking oligos in a single step PCR (table to SI Figure 9)

| Abbrevation in Figure | Plant species |
|---|---|
| *Amar* | *Amaranthus spp.* |
| *A.car* | *Alternantera cardinalis* |
| *A.ses* | *Alternantera sessilis* |
| *A.ten* | *Alternantera tenella* |
| *Pha* | *Phaseolus vulgaris* |
| *Cle_CR* | *Cleome spp.* |
| *C.gyn* | *Cleome gynandra* |
| *F.bid* | *Flaveria bidentis* |
| *F.ram* | *Flaveria ramossisima* |
| *F.rob* | *Flaveria robusta* |
| *F.tri* | *Flaveria trinervia* |
| *Z.mays* | *Zea mays* |
| *At* | *Arabidopsis thaliana* |
| *Arr* | *Arrenatherum elatius* |
| *Bro* | *Bromus erectus* |
| *Gal* | *Galium album* |
| *Ger* | *Geranium pratense* |
| *Lot* | *Lotus corniculatius* |
| *Med* | *Medicago X* |
| *Pla* | *Plantago lanceolata* |
| *Tar* | *Taraxacum officinale* |

**Table S5.** Plants used for testing the universal blocking oligos in a full sequencing run

| Organism Chloroplast | GenBank ID |
|---|---|
| *Bromus erectus* | Collected at Jena Experiment (50°92'77.3"N,11°57'69.7"E), August 2018 |
| *Plantago lanceolate* | Collected at Jena Experiment (50°92'77.3"N,11°57'69.7"E), August 2018 |
| *Lotus corniculatus* | Collected at Jena Experiment (50°92'77.3"N,11°57'69.7"E), August 2018 |
| *Amaranth spec.* | Collected at Fluss Land Jena (50°54'32.9"N 11°34'42.4"E), Summer 2018 |
| *Arabidopsis thaliana Col-0* | Collected from a common garden at Neugasse 23, 07749 Jena, Spring 2019 |

**Table S6.** Composition of the mixed-kingdom mock community.

| Kingdom | Organisms | Rel. Concentration [Even \| Uneven] | Source |
|---|---|---|---|
| Bacteria | *Escherichia coli* | 1 \| 1000 | 1 |
| | *Rhodobacter sphaeroides* | 1 \| 1000 | 1 |
| | *Staphylococcus epidermidis* | 1 \| 1000 | 1 |
| | *Streptococcus mutans* | 1 \| 1000 | 1 |
| | *Bacillus cereus* | 1 \| 100 | 1 |
| | *Clostridium beijerinckii* | 1 \| 100 | 1 |
| | *Pseudomonas aeruginosa* | 1 \| 100 | 1 |
| | *Staphylococcus aureus* | 1 \| 100 | 1 |
| | *Streptococcus agalactiae* | 1 \| 100 | 1 |
| | *Acinetobacter baumanii* | 1 \| 10 | 1 |
| | *Helicobacter pylori* | 1 \| 10 | 1 |
| | *Lactobacillus gasseri* | 1 \| 10 | 1 |
| | *Listeria monocytogenes* | 1 \| 10 | 1 |
| | *Neisseria meningitidis* | 1 \| 10 | 1 |
| | *Propionibacterium acnes* | 1 \| 10 | 1 |
| | *Actinomyces odontolyticus* | 1 \| 1 | 1 |
| | *Bacteroides vulgatus* | 1 \| 1 | 1 |
| | *Deinococcus radiodurans* | 1 \| 1 | 1 |
| | *Enterococcus faecalis* | 1 \| 1 | 1 |
| | *Streptococcus pneumoniae* | 1 \| 1 | 1 |
| Fungi | *Colletotrichum* sp. | 1 \| 100 | 2 |
| | *Galactomyces* sp. | 1 \| 10 | 2 |
| | *Pseudozyma* sp. | 1 \| 1 | 2 |
| | *Rhizopus* sp. | 1 \| 1 | 2 |
| Oomycetes | *Pythium ultimum* | 1 \| 100 | 3 |
| | *Phytophthora capsici* LT1534 | 1 \| 10 | 4 |
| | *Saprolegnia* sp. (1) | 1 \| 1 | 2 |
| | *Saprolegnia* sp. (2) | 1 \| 1 | 2 |

1. BEI Resources product number HM-782D | HM-783D
2. Isolated for this work; The genus was identified via sequencing of the ITS region and a BLAST search against the NCBI database.
3. CBS fungal database strain number CBS 805.95
4. Schornack et al., 2010[1]

**Table S7.** Organisms used for designing primers in the 5.8S rRNA gene region of oomycetes for amplicon sequencing and qPCR quantification.

| Organism | GenBank ID |
|---|---|
| *Achlya bisexualis* CBS10262 | HQ643087.1 |
| *Achlya spinosa* CBS57667 | HQ643109.1 |
| *Albugo candida* AC2V | HQ643111.1 |
| *Albugo hohenheimia* 60A | GU292154.1 |
| *Albugo laibachii* 44A | GU292146.1 |
| *Albugo leimonios* 67A | GU292160.1 |
| *Albugo portulacae* A19 | DQ643921.1 |
| *Albugo tragopogonis* | AF241771.1 |
| *Aphanomyces astaci* FDL458 | AY455772.1 |
| *Aphanomyces cochlioides* CBS47771 | HQ643115.1 |
| *Aphanomyces euteiches* CBS15473 | HQ643119.1 |
| *Basidiophora entospora* H. Voglmayr H.V.122 | EF553487.2 |
| *Bremia lactucae* H. Voglmayr H.V.527 | EF553505.1 |
| *Diasporangium sp.* CCD-2007 | EF198150.1 |
| *Halophytophthora avicenniae* CBS18885 | HQ643147.1 |
| *Halophytophthora epistomium* CBS59085 | HQ643220.1 |
| *Hyaloperonospora brassicae* | JF975613.1 |
| *Hyaloperonospora parasitica* GG158 | EU049283.1 |
| *Leptolegnia caudata* CBS68069 | HQ643137.1 |
| *Perofascia lepidii* SMK 17250 | AY211013.1 |
| *Peronospora belbahrii* PeBGe_HU | HQ702191.1 |
| *Peronospora cristata* BO001 | AY225482.1 |
| *Phytophthora brassicae* CBS782.97 | AF266801.1 |
| *Phytophthora capsici* TARI 9232 | GU111645.1 |
| *Phytophthora infestans* PIT10 | HQ191488.1 |
| *Phytophthora melonis* NN-1 | JN054403.1 |
| *Phytopythium litorale* | JQ898465.1 |
| *Plasmopara halstedii* | AY773346.1 |
| *Plasmopara viticola* | AY742739.1 |
| *Pythiogeton zeae* Lev3132 | HQ643405.1 |
| *Pythiopsis cymosa* CBS 261.34 | DQ393552.1 |
| *Pythium irregulare* | JQ898464.1 |
| *Pythium ultimum* var. sporangiiferum | JQ898477.1 |
| *Pythium vexans* | JQ898479.1 |
| *Saprolegnia diclina* ATCC 90215 | AY455775.1 |
| *Saprolegnia parasitica* NJM9880 | AY455776.1 |
| *Saprolegnia sp.* CAL-2011 rodrigueziana | HQ644006.1 |
| *Saprolegnia* sp. UNCW177 | DQ393512.1 |
| *Wilsoniana amaranthi* FR0046015 | JN849471.1 |

### Supporting Notes

***<u>Testing effects of variations on library preparation and sequencing protocol A</u>***

*Number of blocking cycles*

We tested the effect of altering the library preparation protocol by altering Protocol A (See Supplementary Methods) to have the bulk of PCR cycles as blocking or extension cycles (10 blocking cycles followed by 25 cycles or 25/10). This had only small, non-significant effects with 10/25 giving slightly more accurate results (**Fig S2 and Fig S3**). By comparing to one-step library preparation without blocking oligos, we confirmed previous observations[2] that a two-step amplification with universal followed by concatenated primers tends to generate more reproducible results than a single step with concatenated primers only, although this was only significant in some loci (**Fig S2 and Fig S3**).

*Custom sequencing primers (Protocol A) vs. Illumina sequencing primers (Protocol B)*

Most Illumina amplicon sequencing approaches rely on standard Illumina sequencing primers which bind outside the universal primer-amplified PCR product. This approach first sequences through the universal primers, which can cause low sequence diversity in the first sequenced bases and overall low read quality. To overcome this, large spikes of PhiX genomic library are commonly used and these reads are later discarded[3]. Alternatively, Protocol A adapts that of Caporaso et al.[4] by using custom sequencing primers that are complementary to the universal amplification primers and the adjacent “linker” regions. A key difference is that we combined custom sequencing primers designed for 8 loci so that all regions could be sequenced in parallel. Thus, sequencing begins with bases immediately following the universal primer which are higher diversity than the amplification primer region itself and using multiple loci adds additional diversity. Therefore, a 15% PhiX spike was sufficient to consistently avoid over-clustering. In Protocol B, we use instead the standard Illumina sequencing primers and designed a variable length “offsetting” adapter into the adapters. With this approach we achieved high quality reads with only a 10% PhiX spike. all adpapters and sequences are available in **File S1a**

(Protocol A) and **S1b** (Protocol B). In both cases, sequencing mock communities and comparing the results to expected distributions resulted in satisfactory results (**Fig S2-S4 and Fig S9**).

**Testing potential contamination caused by using universal primers in the blocking cycles**

Since the first PCR step uses non-indexed primers, this approach might increase the risk of cross-contamination. Therefore, we prepared sequencing libraries for three negative controls (two blank DNA extractions and one PCR with water as template) together with 381 plant leaf samples and added the whole amount of negative control library to the sequencing run (**Fig S12**). A maximum of 142 reads were obtained in the three libraries, corresponding to ~2% of the number of reads obtained for *A. thaliana* leaf samples in the same run (average of 6.3 10^3 reads per sample, not shown). There were 362 "contaminant" OTUs with only ~ 13 OTUs exhibiting more than 10 total reads. The most abundant ones were classified as the water contaminant bacterium "*Pelomonas puraquae*" (~48 total reads) and "*Pseudomonas*" (~26 total reads).

### Supporting Methods

***Mixed kingdom mock community preparation***

To optimize the multi-locus method, we used a mock community with known relative abundances of bacteria, fungi and oomycetes. Bacterial mixes (even and staggered concentrations) were ordered from the Biodefense and Emerging Infections Research Resources Repository (BEI) (**Table S6**). Fungal and oomycete communities were prepared, each using quantified genomic DNA extractions of four microorganisms either from the CBS-KNAW Fungal Biodiversity Centre or which were isolated for this study. In short, we isolated genomic DNA from select cultures grown on potato dextrose agar using bead beating in SDS lysis buffer with lysozyme and proteinase K, RNAse treatment, and phenol/chloroform extraction followed by an additional cleanup with 20% Chelex-100. We relatively quantified the amplifiable ITS units in DNA extracts from each organism with qPCR using fungal (FR1/FF390 from[5]) and oomycete (designed for this study - see **Supporting Methods** below) ITS primers. Next, we combined the extracted DNA of fungi or oomycetes plus the bacterial mixes to end up with even or uneven amounts of amplifiable ITS from each organism (**Table S6**). To test effects of host contamination and non-target amplification, we combined mock communities that were 5% microbial and 95% genomic DNA extracted as above from axenically grown *A. thaliana* Ws-0 based on PicoGreen (Life Technologies, Inc) quantification.

***Library preparation and multi-locus amplicon sequencing (Protocol A)***

For each sample template, all targeted regions were amplified in parallel on a single PCR plate and this was performed in triplicate. Reactions contained 0.2 µL Q5 high-fidelity DNA polymerase (New England Biolabs, Inc), 0.5 µL template, 1X Q5 GC Buffer, 1x Q5 5x reaction buffer, 0.08 µM each forward and reverse primer (**Table S1** and **File S1**), 0.25 µM of each blocking oligonucleotide (**File S1**) or nuclease free water, 225 µM dNTP and 10.33 µL nuclease free water. In the first step, triplicate plates were run in parallel on three independent thermocycler blocks (Bio-Rad Laboratories, Inc.) at 95 °C for 40 sec, followed by a defined number of cycles of 95 °C for 35 sec, 55 °C for 45 sec, 72 °C for 15 sec with a final elongation at 72 °C for 3 min. The three replicate reactions were then combined. For reactions that would be subject to a second PCR step, 10 µL was recovered for an enzymatic cleanup with Antarctic phosphatase and Exonuclease I (New England Biolabs, Inc) to degrade leftover primers and inactivate nucleotides (0.5 µL each enzyme with 1.22 µL Antarctic phosphatase buffer at 37 °C for 30 minutes followed by 80 °C for 15 min). The products of the enzymatic cleanup reaction were then used as template in a second, 50 µL PCR reaction with an identical setup as in the first step. For the 11 templates, we used 11 different reverse primers that were identical for each group of amplified microbes except for the 12-bp index[4] that would be used later to identify sequencing products in combined libraries (**File S1a**). Final products were purified using 0.8x volume Ampure XP purification beads (Beckman-Coulter, Inc) and eluted in 40 µL according to the manufacturers instructions.

Amplified and cleaned gene products were fluorescently quantified with PicoGreen (Life Technologies, Inc) using salmon sperm DNA (Invitrogen, Inc) as standard in two independent measurements at both 100x and 1000x dilutions. The results were averaged, and molar DNA concentrations were calculated using the approximate length of amplicons of each region while assuming a 500 bp average length for the salmon sperm DNA. The quantified amplicon libraries from all samples and regions were combined in equimolar concentrations and concentrated 10x using 0.8x volume Ampure XP purification beads before quantification on a Qubit (Thermo Fisher Scientific, Inc).

The library was loaded onto a MiSeq lane spiked with 15% PhiX genomic DNA to ensure high enough sequence diversity. We adapted the custom sequencing primer approach of Caporaso et al. [4] to multiple loci (**Fig S1C** and **File S1a**). 3.4 µL of a 100 pM solution of each of the custom sequencing primers (**Fig S1C** and **File S1a**) were combined with the standard Illumina sequencing primers for each of the forward, reverse, and index sequencing reactions. Sequencing was performed for 500 cycles to recover 250 bp of information in the forward and reverse directions (**Fig S1C**).

***<u>Processing multi-locus amplicon sequencing data with a multi-step OTU picking approach (Protocol A)</u>***

We used a custom data processing pipeline to handle raw multi-locus read data (further details for each of the steps can be found in[6]). The data from mock community amplification and from *A. thaliana* leaf microbiome amplification were processed separately from one another. In short, the pipeline first demultiplexed and quality filtered data using QIIME 1.8.0[7] and divided results into the amplified target regions. Next, we trimmed adapter sequences from distal read ends using Cutadapt 1.2.1[8]. We merged the remaining paired reads using Pandaseq[9]. At this point in the pipeline, we had for each of the microbial groups aligned reads, unaligned reads (those that could not be merged because of a long amplicon or strong quality trimming), as well as read 1 and read 2 orphans from the first quality filtering step. For each of the 6 or 8 groups we concatenated all reads together (aligned, unaligned, and orphans). For fungal and oomycete ITS datasets, we used ITSx 1.0.8[10] to trim reads to only the ITS region. This was not possible for the oomycete ITS1 region because neighboring rRNA gene regions were too small for detection. Next, for all groups, we performed denovo and reference-based chimera checking with USEARCH 6.1[11] using the greengenes 13_8 release[12] or our custom eukaryote databases (see below) as a reference for bacteria and fungi/oomcyetes, respectively. In order to group orphan and paired reads which differ in length, we developed a multi-step OTU picking protocol. In short, we first picked "prefix" OTUs then from these picked "suffix" OTUs using the prefix/suffix OTU picker in QIIME. Finally, for all 6 groups we picked final OTUs using UCLUST at 97% identity and picked representative sequences for each OTU. In the final OTU map, we removed OTUs with < 2 reads.

We assigned taxonomy to the final set of OTUs using the RDP taxonomy assigner[13] implemented in QIIME at a confidence level of 0.8 for 16S sequences (against the Greengenes 16S rRNA database release 13_8) and 0.65 for ITS and 0.8 for 18S sequences (against custom eukaryote ITS and 18S databases, respectively – see **Supporting Methods** below). We used QIIME to generate an OTU table and to filter OTUs present in < 2 samples and with < 50 reads. As a final filtering step, we filtered "Unclassified" or "Unassigned" reads from the 16S and 18S OTU tables. From the ITS OTU tables we filtered "Unassignable", "Unclassified" or the kingdom "unidentified". Because our ITS database contained more than one eukaryotic kingdom, we were able to identify significant amounts of non-target reads in some samples (e.g., oomycetes in fungal ITS libraries). Therefore, we filtered fungi from oomycete ITS OTU tables and oomycetes from fungal ITS OTU tables. For analyses where host-derived OTUs should not be included, we filtered any OTU in the class “Chloroplast” or the order “Rickettsiales” (primarily host mitochondria 16S) from the 16S OTU tables and reads in the kingdom “Viridiplantae” from the ITS OTU tables.

We summarized bacterial, fungal and oomycete OTU tables by taxonomic rank, converted abundances to relative values and plotted the genus-level taxonomic distribution directly from this data with the package ggplots2 in R. To calculate distances of samples from expected, we added the expected distributions (**Table 1**) to the OTU tables and summarized taxa at the family level. After removing “host“-derived reads, we calculated Bray-Curtis distances between samples using Vegan[14]. We plotted distances from expected distributions in boxplots using ggplots2. Each box represents three “replicate” libraries generated with the same mixed kingdom mock community template but with differing amounts of “host“ DNA added.

Scripts used to generate OTU tables from the raw data, as well as OTU tables and metadata files to recreate the main figures are being made publicly available on Figshare (https://figshare.com/projects/Obtaining_deeper_insights_into_microbiome_diversity_using_a_simple_method_to_block_host_and_non-targets_in_amplicon_sequencing_/89504).

***DNA extraction of wild leaf samples for multi-plant sequencing***

Three individual leaves were collected from the species *Plantago lanceolata*, *Lotus corniculatus*, *Bromus erectus*, were rinsed with autoclaved water and immediately frozen on dry ice. They were stored at -80°C until DNA extraction. For extracting DNA, we ground the leaves while frozen in liquid nitrogen. A small amount of ground leaf material was added to screw-cap tubes and was bead beat with 0.2g of 0.25-0.5mm glass beads (30s at 1400rpm on a BioSpec mini bead beater 96). 200uL CTAB extraction buffer (100mM TRIS pH 8.0, 20mM EDTA pH 8.0, 1.5 M NaCl, 2% cetylimethylammoniumbromid, 1%, polyvinylpyrrolidone (MW 40,000) in nuclease-free water) was added incubated at 37C for 10 min. Next, the solution was spun at 20,000g at 4C for 5 min and supernatant transferred to a new tube. DNA was precipitated with 200uL cold 100% ethanol and centrifuging at 13000rpm at 4C for 5 min. The pellet was rinsed in 70% ethanol, allowed to dry and resuspended in 100uL TRIS-HCl buffer (pH 8). This was followed by a short clean up using home-made Sera-Mag purification beads[15]. For *Arabdiopsis thaliana* Col-0 and *Amaranthus spec*. leaves, the protocol was similar, except that the plant material was not crushed with liquid nitrogen: The plant material was added to the 2mL screw-cap tubes which in addition to the glass beads also contained two metal beads to crush the material. The sample underwent beat-beading right after it left the freezer to optimize the crushing process.

***Library preparation for multi-plant amplicon sequencing (Protocol B)***

*Blocking cycles*

For each sample template, all targeted regions were amplified in parallel on the same 384-well PCR plate with each reaction in quadruplicate. First, 40uL reactions were prepared with 0.8 µL Kapa taq DNA polymerase (KAPA Biosystems), 1 µL genomic DNA template, 1X Kapa GC Buffer, 10 µM of each forward and reverse primer (**Table S1** and **File S1**), 33.33 µM of each blocking oligonucleotide (**File S1**) or nuclease free water, 10 mM dNTP and 26.4 µL nuclease free water. The mixes were divided into four 10uL reactions with a 96-well pipetter (Platemaster, Gilson) on to a 384-well plate for independent amplificaiton. These were run on a thermocycler block at 95 °C for 3 min, followed by a 10 or 15 cycles of 98 °C for 20 sec, 55 °C for 30 sec, 72 °C for 15 sec or for 40 sec (10 cycles for 15 sec, 15 cycles for 40 sec) with a final elongation at 72 °C for 2 min.

*Extension cycles*

The four replicate reactions were re-combined and 10 µL was recovered for an enzymatic cleanup with Antarctic phosphatase and Exonuclease I (New England Biolabs, Inc) to degrade leftover primers and inactivate nucleotides (0.5 µL each enzyme with 1.22 µL Antarctic phosphatase buffer at 37 °C for 30 minutes followed by 80 °C for 15 min). The products of the enzymatic cleanup reaction were then used as template in a second, 15 µL PCR reaction with an identical setup as in the first step and for 25 or 20 cycles. The only difference was the use of concatenated barcoded primers that each had with a 12-bp index (sequences in **File S1**).

*Library normalization*

Final products were purified using 0.8x volume home-made Sera-Mag purification beads and eluted in 40 µL TRIS-HCl. Amplified and cleaned gene products were fluorescently quantified on an Agarose gel using Roti Stain. Lanes on the gel were quantified via ImageJ, combined in roughly equimolar concentrations and concentrated again with 0.8x volume Sera-Mag purification beads. The exact length of the amplicons was analyzed on a 2100 Bioanalyzer Instrument and the library was quantified on a Qubit (Thermo Fisher Scientific, Inc). The library was denatured and then loaded onto a MiSeq lane spiked with 10% PhiX genomic DNA to ensure high enough sequence diversity. Sequencing was performed for 500 cycles to recover 250 bp of information in the forward and reverse directions with standard Illumina sequencing primers.

***<u>Processing multi-plant amplicon sequencing data (Protocol B)</u>***

We split the amplicon sequencing data on indices and trimmed the adapter sequences from distal read ends using Cutadapt 1.2.1[8]. We then clustered amplicon sequencing data into amplicon sequencing variants "ASVs" using dada2 by first filtering them based on their quality (truncLen=c(200,200)). Using dada2 we then dereplicated the sequences to eliminate exactly redundant reads. Data was denoised using the error rate information and calls amplicon sequence variants (ASVs) in the forward and reverse reads. We merged the forward and reverse reads by keeping data with overlapping regions, discarding non-overlaps and concatenating the rest. We removed chimeric sequences and retrieved a sequence table from the merged data. We assigned taxonomy to the final set of ASVs using the Silva[16] database (16S: “silva_nr_v132_train_set.fa” downloaded at 19.03.2019 and 18S: “silva_132.18s.99_rep_set.dada2.fa” downloaded at 16.01.2020). We performed downstream analyzes in R with phyloseq. For analyses where host-derived ASVs should not be included, we filtered any ASV in the class “Chloroplast” from the 16S ASV tables and from the class “Chloroplastida” from the 18S ASV tables. Scripts used to generate ASV tables from the raw data, as well as ASV tables and metadata files to recreate the main figures are being made publicly available on Figshare (https://figshare.com/projects/Obtaining_deeper_insights_into_microbiome_diversity_using_a_simple_method_to_block_host_and_non-targets_in_amplicon_sequencing_/89504).

***Primer design for oomycete amplification and quantification***

To relatively quantify ITS units in oomycete species and to design additional primers to make sequencing of the oomycete ITS1 and ITS2 regions possible, we designed primers in the oomycete 5.8S rRNA gene region as follows. First, we downloaded nearly complete ITS regions (most include a partial 18S rRNA gene, ITS region 1, 5.8s rRNA gene, ITS region 2, and partial 28S rRNA gene) for 39 oomycete species that are widespread across the oomycete phylogeny (**Table S3**). We aligned the sequences using the SeaView 4.3.5[17] implementation of MUSCLE[18]. After alignment, we identified a highly conserved region in the 5.8S gene starting at position 748. For qPCR, we designed primers to amplify a 91 bp portion of this region. The forward primer (5.8sF) includes positions 748 – 768 of the alignment: ACTTTCAGCAGTGGATGTCTA. The reverse primer (5.8sR) was designed complimentary to positions 819 – 839: GATGACTCACTGAATTCTGCA. For amplicon sequencing, we designed a reverse primer near the 5' end of the 5.8S gene to amplify the ITS1 region with the ITS1-O forward primer. This primer (5.8S-O-R) is complementary to positions 753-772: AGCCTAGACATCCACTGCTG. We also designed a forward primer near the 3' end of the same gene to amplify the ITS2 region with the ITS7 reverse primer. Primer 5.8S-O-F-deg is complementary to positions 846-864: TTGAACGCAYATTGCACTT.

***Eukaryote ITS and 18S database creation***

*ITS database customization*

To enable taxonomy assignment for oomycete ITS reads, we needed to address the lack of oomycete taxa coverage in fungal ITS databases. Further, conserved regions where ITS primers are designed have some similarity to non-targeted groups, so a broader database is needed to identify non-target amplicons, e.g., amplicons from oomycetes and host plants in fungal ITS amplicon libraries. To address this problem, we created a more diverse eukaryote ITS reference database for use in processing ITS reads. To do so, we combined the UNITE fungal ITS database ([19], QIIME-compatible version, downloaded on July 7th, 2014), the *Arabidopsis thaliana* ITS sequence, and a custom made oomycete ITS database. To create the oomcyete database, we wrote a script to recover all nucleotide sequences from NCBI corresponding to taxid 4762 (oomycetes). For each recovered sequence, we then used the GIDs to recover the corresponding taxonomy and created a 6-level taxonomy file (in a QIIME 1.8.0-compatible format similar to the Greengenes 16S rRNA and UNITE ITS databases). With the oomycete sequences, we picked OTUs using the QIIME implementation of UCLUST with default settings at a 97% sequence similarity. To clean up the database, we located sequences that contained at least the 5.8s region (reads containing ITS1, 5.8S rRNA gene and ITS2 were nearly full length and most useful) by searching OTUs with BLAST[20] against a database of fungal 5.8S rRNA sequences, allowing poor matches (e-value 100). We discarded reads without a 5.8s region, only kept sequences that were at least 80bp long and discarding reads containing degenerate bases. We then trimmed the previously made taxonomy files to those sequences that were remaining after all processing steps. With the combined fungal, A. thaliana, and oomycete databases, we used ITSx 1.0.8[10] to divide the database into ITS1 and ITS2 regions which were used as databases in further analyses.

*PR2 database customization*

Similarly to the ITS database, we created PR2 databases for the V4 and V9 regions of the 18S rRNA genes. We used a custom script to pick out the V4 and V9 regions based on primer sequences for the regions from the entire PR2 database[21]. Essentially, the script used BLAST against the PR2 databse to locate where the universal primers for each region (**File S1a**) match. Based on the results, the database was then divided into the two regions. Next we cleaned the database by removing sequences with empty headers and sequences shorter than 80 bp as well as sequences with degenerate bases. We then picked OTUs and representative sequences exactly as for the oomycete ITS sequences above. We then recovered the PR2 taxonomy and created a taxonomy file in 6 levels compatible with QIIME, then trimmed it to match the remaining sequences after the filtering and OTU picking steps.

All databases are being made publicly available on Figshare: https://figshare.com/projects/Obtaining_deeper_insights_into_microbiome_diversity_using_a_simple_method_to_block_host_and_non-targets_in_amplicon_sequencing_/89504
